# Spatial transcriptomics implicates the thalamus and cortex in autism and schizophrenia

**DOI:** 10.64898/2026.05.15.724759

**Authors:** David M. Young, Ruchira Sharma, Narjes Rohani, Chimmi Dema, Lindsay Liang, Bernie Devlin, Devanand S. Manoli, Stephan J. Sanders

## Abstract

The past decade has seen tremendous progress in the identification of genes associated with complex neuropsychiatric disorders, including autism spectrum disorder (ASD) and schizophrenia. Expression patterns of these genes in single cell data strongly implicate excitatory and inhibitory neurons; however, there are limited data on the brain regions involved - a critical question for neurobiology. Spatial transcriptomics provide an opportunity to perform systematic multiregional analyses to provide insights into this question. Here, we have generated a spatial transcriptomics dataset encompassing the diverse anatomical territories of the adult mouse brain sagittal midsection. We compare neuropsychiatric gene enrichment by applying Gene Fraction Enrichment Score (GFES), a novel statistic method that controls for differing neuronal proportions across regions. ASD-associated genes identified by exome sequencing were most enriched in the thalamus followed by the cortex. Schizophrenia genes from genome-wide association studies were also enriched in the thalamus, along with the hippocampus and cortex. These findings add to the evidence that the thalamus plays a major role in neuropsychiatric disorders whilst supporting roles for the cortex and hippocampus. The results highlight shared and distinct patterns for pleiotropic brain disorders that could elucidate common underlying mechanisms and circuitry.

## Introduction

Some brain-related disorders are caused by specific lesions in defined brain regions, such as the substantia nigra in Parkinson’s Disease or the internal capsule in hemiplegic strokes. For many others, including autism spectrum disorder (ASD) and schizophrenia the brain regions involved remain poorly defined. Resolving the brain regions involved is of critical importance to identify molecular, cellular, or circuit level etiology and for translation, including biodistribution and neuromodulation.

In humans, neuroimaging offers a practical route to systematic analyses across multiple regions of the living brain, providing insights into structure, circuitry, and function. Analysis of large cohorts have identified subtle diffuse findings. These include structural changes, such as increased cortical thickness in ASD and decreased in schizophrenia[1–3] and functional changes, including disruption of the default mode network, often with decreased connectivity[3, 4]. The default mode network includes regions of the cortex and the hippocampus. Interpretation of neuroimaging data is complicated by small effect sizes and substantial variation among individuals and across studies.

Both ASD and schizophrenia are highly heritable, supporting a major role for genetic factors. Analyses of DNA from large cohorts of ASD and schizophrenia families, cases, and controls have yielded lists of genes associated with both disorders[5– 10]. In both disorders these disorder-associated genes represent a foundation of causal factors from which to investigate downstream neurobiology. Behavioral studies of animal models of individual genes have yielded mixed results, implicating most major brain regions[11]. In contrast, assessing the enrichment of these ASD and schizophrenia gene lists at the RNA level with transcriptomic data has clearly implicated excitatory and, to a lesser extent in ASD, inhibitory neurons in both disorders[6, 8, 12–14]. Most of the genes are expressed from the midfetal period onwards[14, 15]. Transcriptional changes can also be assessed in *postmortem* case/control analyses and have yielded altered expression of neuronal genes in the cortex from analyses of bulk tissue[16]. To date, other brain regions have not been assessed in sufficiently powered samples.

Ideally, we could identify brain regions where disorder-associated genes are commonly expressed. Spatial transcriptomics has emerged as a systematic method built upon the next-generation sequencing methods of scRNA-seq to profile the full transcriptome in a spatial context by simultaneously recording the position and identity of each RNA transcript[17]. Spatial transcriptomics approaches[17, 18, 19] have thus far been applied to a wide variety of brain disorders, including Alzheimer’s disease[20, 21] to amyotrophic lateral sclerosis (ALS)[22]. By measuring across the transcriptome without prior selection of transcripts, spatial transcriptomics highlights regions enriched for disorder-associated genes, pointing to where they are more likely to be active and involved in disease when dysregulated.

Here, we present a multiregional, cross-disorder, spatial transcriptomics analysis of gene convergence in the adult mouse brain. We selected the sagittal midsection of the brain for its wide diversity of anatomical and functional territories and identified 29 clusters, which largely correspond to anatomical regions and sub-regions. To study the convergent expression of risk gene sets for polygenic neuropsychiatric disorders, we introduce a novel gene enrichment metric (GFES) to highlight spots expressing a preponderance of genes associated with each disorder whilst correcting for cell type proportion. Applying this to ASD and schizophrenia we observed enrichment in the murine thalamus and olfactory bulb, alongside the cortex and hippocampus.

## Results

### Samples assayed from the sagittal midsection of mature mouse brain

We elected to assay sagittal midsections of the mature mouse brain (postnatal day 56 C57BL/6J; three female, one male) to enable multiple major brain regions to be assessed systematically, including cortex, hippocampus, basal ganglia, thalamus, hypothalamus, and midbrain (Fig. 1a). Each 14µm section spanned the full extent of the brain dorsoventrally and from the posterior olfactory area to the midbrain anteroposteriorly, covering a substantial fraction of the brain’s rostral-caudal and inferior-superior axes. Four sagittal sections were assessed from each mouse brain separated by about 100µm to capture a degree of medial-lateral variability (Fig. 1b-c).

**Figure 1:**
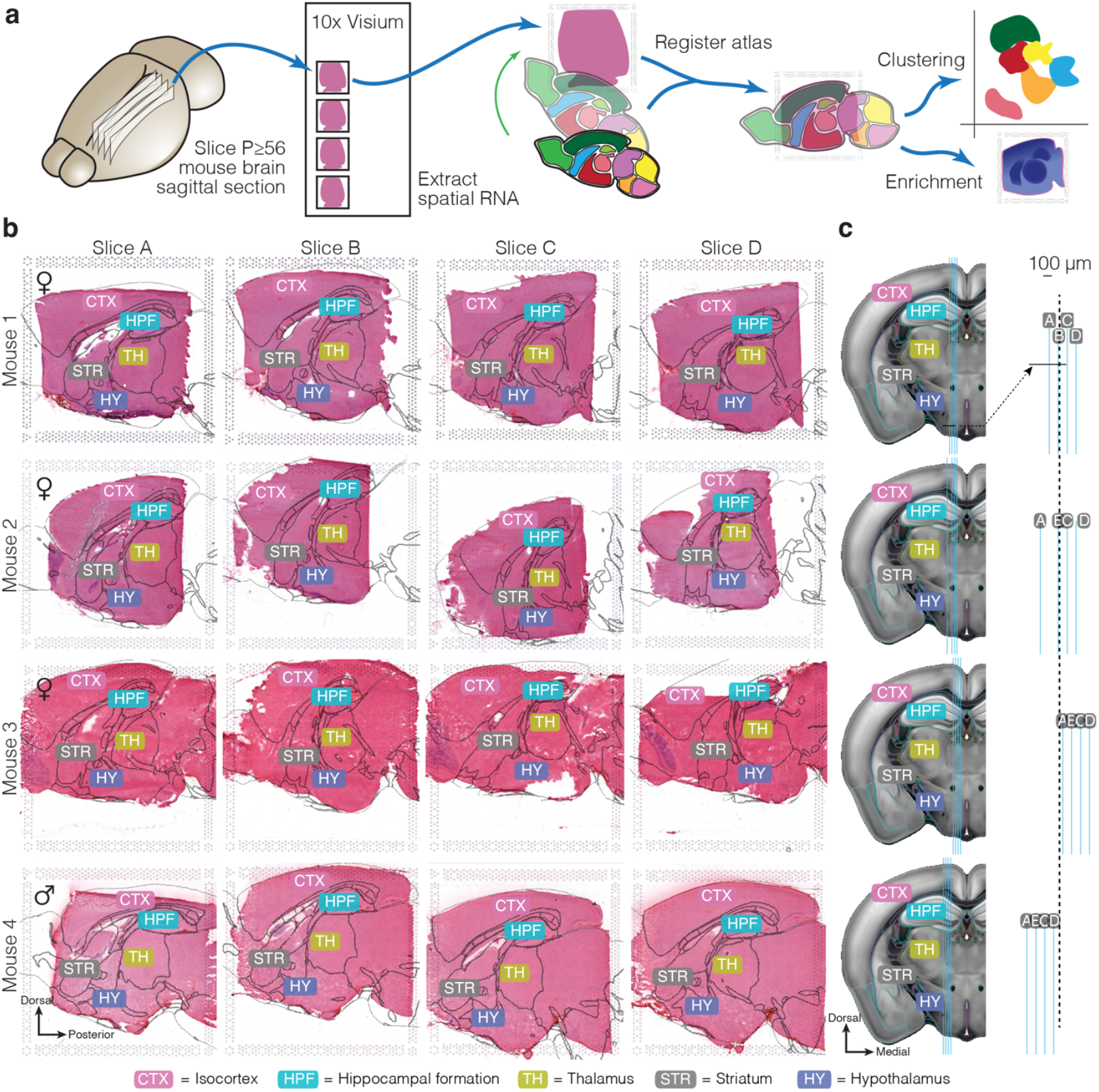
Experimental overview and anatomical orientation. **a)** Serial sagittal adult mouse brain sections were collected onto 10x Visium capture areas, which extracted RNA onto spatially barcoded spots. Tissue sections were segmented by atlas registration to identify clusters and enriched regions. **b)** Tissue samples from 16 sections in four mouse brains (rows) by serial slice (column) from lateral (left) to medial (right). Registered anatomical boundaries are overlaid and labelled. **c)** Corresponding slice positions represented in the coronal plane (left) and magnified for relative positioning (right).

Spatial gene expression was assayed using the 10x Visium platform. All four fresh frozen sections from each mouse were placed on a single Visium slide, which contains four capture areas of 6.5 × 6.5mm. Each capture area contained 4,992 spots of 55µm diameter arrayed in a grid-like pattern with 100µm between spot centers. Each spot captures RNA transcripts lysed from the overlying tissue, including a unique spatial barcode to identify the location of each transcript. An image of the tissue section was captured using a hematoxylin & eosin (H&E) stain and whole slide imaging microscope.

### Anatomical alignment

Histological images were registered to the well-validated Allen Common Coordinate Framework version 3 (CCFv3) atlas[23] that represents an age-appropriate (P56) reference with 3D labels appropriate for all orthogonal axes, including the sagittal orientation. To identify the anatomical location of each slice on the medial-lateral axis, each microscopy image was compared to sagittal planes from the CCFv3 to identify the best match (Fig. 1c). Both rigid and deformable registration of each atlas plane to the corresponding tissue section was performed using the Elastix toolkit as implemented in our MagellanMapper software[24, 25], thereby assigning an anatomical structure to each spot (Fig. S1).

### Transcriptomic quality control and clustering

Sequencing data were aligned to the mm10 mouse genome using STAR[26]. Data were processed using 10x Genomics’ Space Ranger to yield tables of Unique Molecular Identifiers (UMIs) per feature per spot and analyzed by Seurat[27]. High quality spots were identified based on feature counts (≥100) and UMI counts (≥100) with thresholds determined by Density-Based Spatial Clustering and Application with Noise (DBSCAN) analysis[28] (Fig. S2a) and under 30% mitochondrial genes. A median of 3,274 high quality, tissue-covered spots were included per section (53,841 total), with a median of 16,541 UMIs per spot, and 4,974 features per spot (Fig. S2b). Seurat’s SCTransform integration pathway was used to integrate and normalize counts across the 16 sections[27].

Applying Louvain clustering[29] to the 16 integrated sections identified 29 clusters (Fig. 2a), largely corresponding to anatomical structures, consistent with expectation[30] (Fig. 2b). Some major structures were subdivided into smaller clusters by layer (e.g., the cortex), by laterality (e.g., midline vs. medial clusters of L4-6 cortex and the thalamus), or by substructure (e.g., ventromedial hypothalamus). Three clusters were more dispersed (CTX-endo, TT-endo, BS-endo), with scattered spots enriched in hemoglobin genes probably representing spots with higher densities of vasculature.

**Figure 2:**
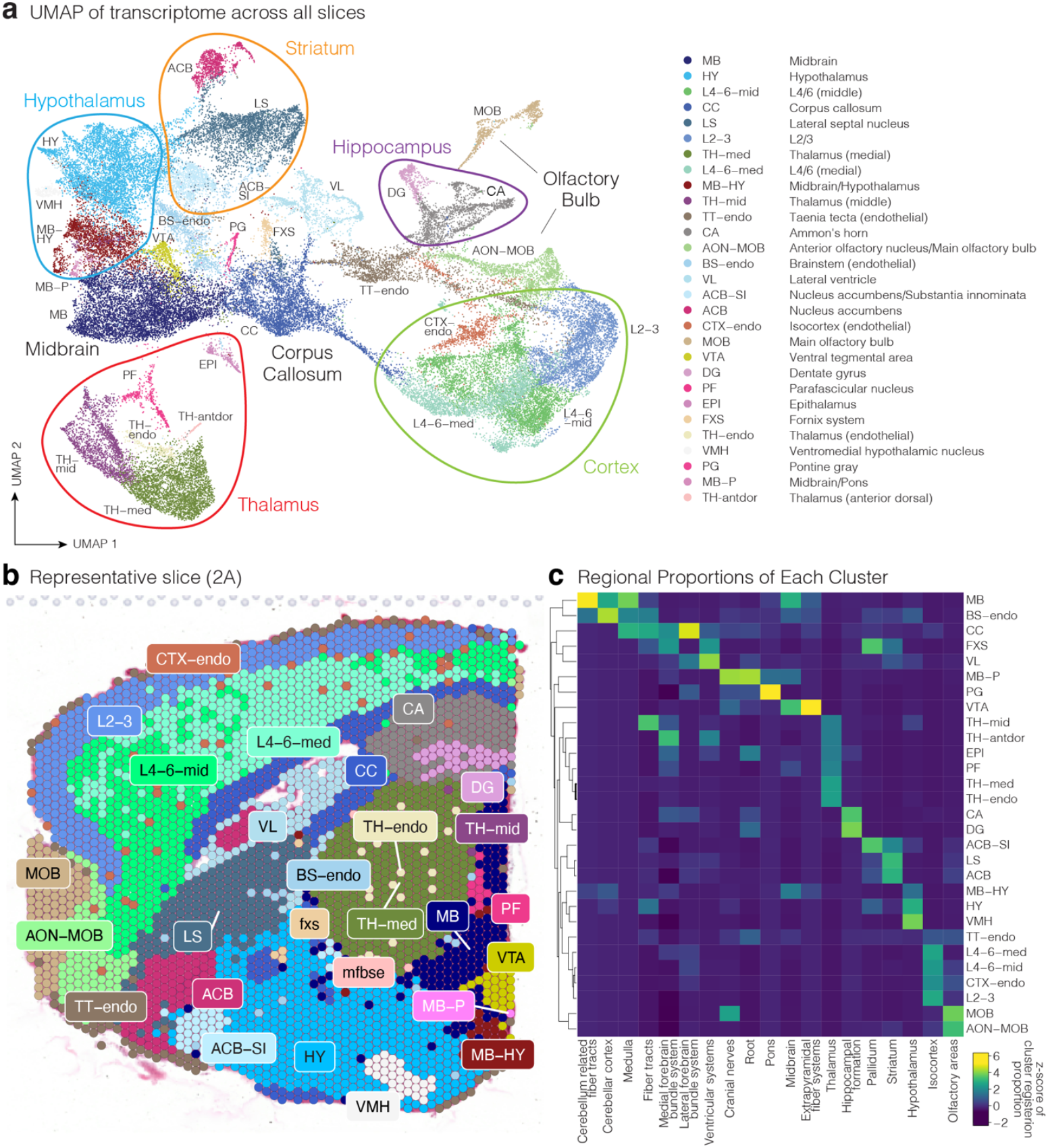
Cluster identification by anatomical alignment. **a)** UMAP of transcriptomic spots with spots are colored by cluster identity. Clusters related to similar anatomical regions are circled and/or labelled. **b)** Spatial clustering shown on a representative slice (Slice 4A, see Fig. 1b). Most clusters aggregated into anatomically distinct groups, though some clusters span multiple regions (e.g., MB-HY) or are dispersed within a region (e.g., TH-endo). **c)** Heatmap of clusters (y-axis) by registered CCFv3 atlas region showing the standardized z-scores of the proportion of each cluster occupied by CCFv3 registered atlas regions (x-axis). Most clusters predominantly match only one region.

### Integrating anatomical registration and transcriptomic clusters

While anatomical registration aligned well to the outer tissue borders, some internal structures remained misaligned. Therefore, we coupled image registration with the transcriptomic data by leveraging ‘anchor’ regions with a clear match between clusters and anatomical structures, such as the thalamus. The centers of these anchor clusters and their corresponding atlas structures were linked to enable cluster-guided registration (Fig. S1). For each cluster, we plotted the proportion of spots assigned to each atlas-defined anatomical region as a heatmap (Fig. 2c). Most clusters matched a single anatomical region, highlighting the anatomical identity of each cluster.

### Cluster marker identification

To further define and validate each cluster, we used Seurat to identify marker genes enriched in specific clusters (Fig. S3), refined the list to genes with large differences in percent expression across clusters, and applied a graded tissue-specificity metric, tau[31, 32]. Many of the resulting markers were substantially enriched in single clusters only (Fig. 3a), providing utility as anatomical markers, despite being less highly ranked based on mean differential fold change across clusters. For example, the marker with the most specificity to the hypothalamus was thyrotropin-releasing hormone, *Trh*, a known marker of hypothalamic neurons, that had been ranked eighth based on differential fold change. Similarly, the highest ranked marker of the L4-6-mid cortex cluster was *Cort*, a gene expressed in cortical inhibitory neurons, whereas the highest differential marker was *Sst*, which happens to be structurally similar to *Cort* but also expressed in the ventromedial nucleus of the hypothalamus (VMH).

**Figure 3:**
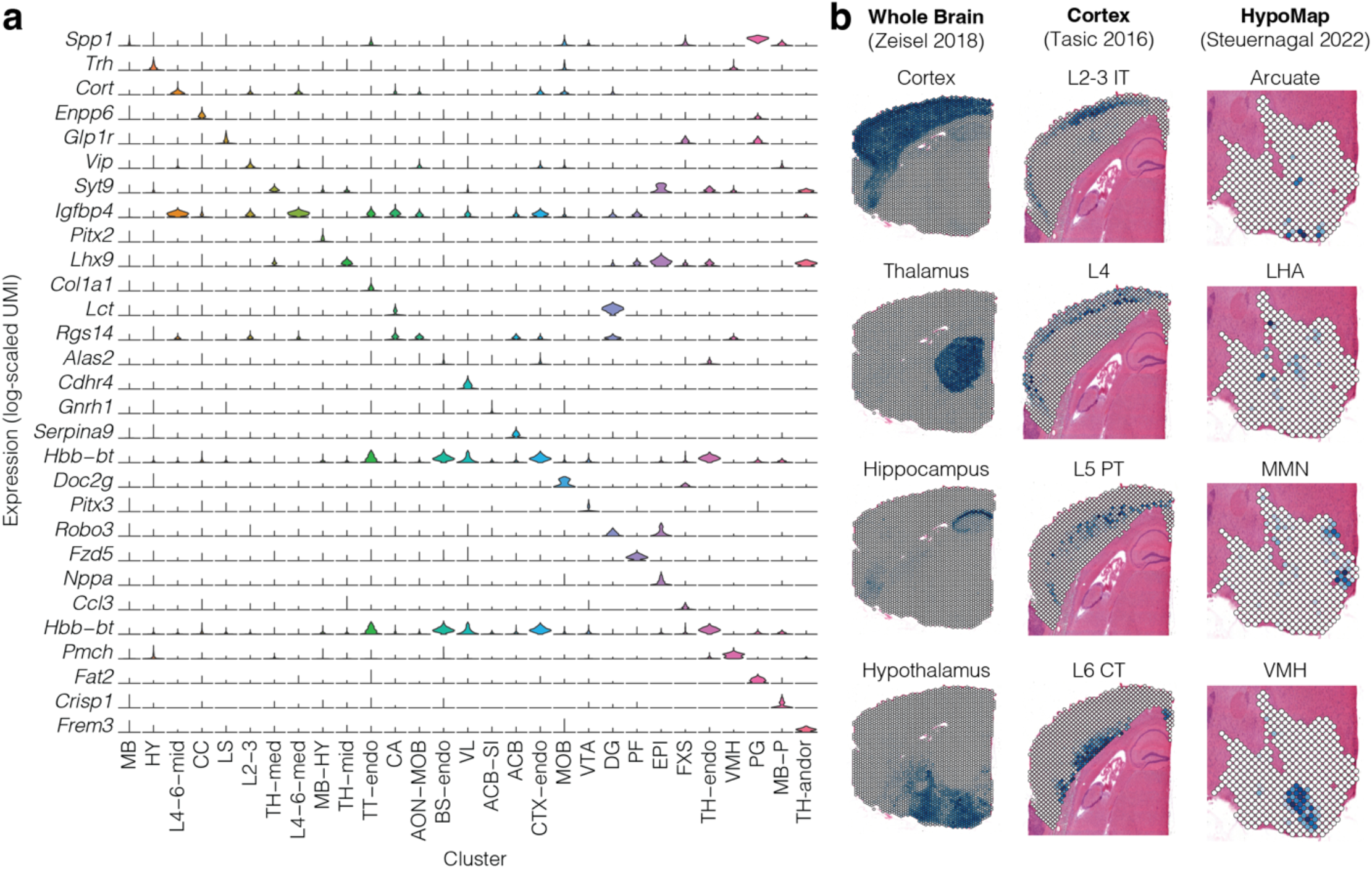
Transcriptomic clusters delineated by tissue-specific markers and deconvolution. **a)** Violin plots show expression levels by cluster of tissue-specific genes identified by cluster-specific differential expression and high tau values. **b)** Cluster anatomical assignments compared to external reference datasets of the whole brain[33], cortex[34], and hypothalamus[35]. For each anatomical spot, Seurat prediction scores are represented by shade, with dark blue representing higher prediction scores, normalized to each spatial slice.

Each spatial spot assays the transcriptome of multiple overlaying cells; deconvolution can estimate the proportion of these cell types through comparison to the transcriptome of known cell types. We used established single-cell/nuclei RNA-seq datasets from specific brain regions (Fig. 3b) to perform deconvolution with RCTD[30, 36] to validate the anatomical assignment of clusters. The Zeisel *et al*. dataset[33] details the transcriptome of over half a million cells from contiguous regions dissected from the nervous system of adolescent mice (P20-30, CD-1/Swiss). Using cells from specific regions (e.g., cortex, thalamus, hippocampus, hypothalamus) as the input for deconvolution, we observed high concordance for spots assigned to the corresponding anatomical cluster (Fig. 3b). Thus, we demonstrate that our clusters are accurately assigned via image registration (Fig. 2), transcriptomic markers (Fig. 3a), and comparison to single cell data from dissected brain regions (Fig. 3b).

Other single-cell/nuclei RNA-seq datasets provide greater resolution within anatomical structures. Considering over 23,000 cells from the neocortex of P53-59 C57BL/6J mice in the Tasic et al. 2016 dataset[34], we observed concordance with excitatory neurons in specific cortical layers matching our anatomical clusters (Fig. 3b). Similarly, integration of 17 datasets to yield a murine hypothalamic atlas for 384,000 cells (HypoMap)[35], enabled specific nuclei of the hypothalamus to be identified in our spatial transcriptomic data (Fig. 3b).

### Spatial gene enrichment corrected for cell type proportion

Having registered the spatial images and defined and validated spatial clusters, we assessed gene enrichment within spatial clusters to yield insights into brain regions related to ASD and schizophrenia. Prior analyses have demonstrated that genes associated with brain phenotypes are enriched in neurons[37–39]. Since the proportion of neurons varies across brain regions (Fig. S4), we must correct for the neuronal percentage per spot to assess regional enrichment. Therefore, we used the Zeisel *et al*. whole-brain dataset[33] to deconvolve each spot to estimate neuronal fraction. As expected, spots in the cortex and hippocampus had high neuronal fractions, while spots in the corpus callosum had low neuronal fractions.

Based on the logic that a gene needs to be expressed to contribute to a disorder, gene enrichment studies often use over-representation analysis to assess whether a defined list of genes are expressed more than expected in a specific cell type[40]. Each gene is defined as expressed or not expressed, based on exceeding a specific threshold, e.g., present in 25% of cells in a cell-type cluster[8]. The fraction of genes on the list that are expressed in the cluster is compared to the total number of genes assessed to calculate a p-value (e.g., Fisher’s exact test). This approach benefits from simplicity; however, the inclusion of covariates, such as neuronal proportion, is complicated, varying the threshold defining ‘expressed’ can alter results, and the total number of expressed genes varies between clusters, confounding interpretation. Of note, the use of a threshold of cells with counts to define gene expression suggests that single cell gene enrichment analysis is measuring whether a gene is highly expressed, rather than an absolute measure of expressed or not expressed. This approach is embraced in Functional Class Storing (FCS) method[41] that uses quantitative expression to assess enrichment, typically generating a score per gene.

To apply gene enrichment analysis to spatial transcriptomic data, we assessed the proportion of total UMIs that aligned to genes in the list per spot, yielding a graded, self-normalized enrichment metric. To compare gene enrichment across clusters, we applied ANCOVA to the per spot metrics, treating clusters as the independent, categorical variable and per spot neuronal proportion as a covariate. This ANCOVA approach calculates regression estimates per cluster that can be compared between clusters. Because each spot is likely to be next to a spot in the same cluster, each spot is not independent, therefore the ANCOVA p-values are not informative. To overcome this limitation, we used a bootstrap procedure in which we assign a subset of spots randomly to each cluster and use ANCOVA to yield regression estimates. With multiple bootstrapping iterations we can define a 95% confidence interval per cluster, enabling significant differences between clusters to be evaluated (Fig. S5). We have termed this graded, self-normalizing, covariate compatible, threshold-free, cell/spot-level metric ‘Gene Fraction Enrichment Score’ (GFES). Through simulation, we demonstrate that GFES can accurately and quantitatively distinguish different patterns of enrichment across transcriptomic clusters (Fig. S6).

### Autism genes are enriched in the thalamus, olfactory bulb, and cortex

We next considered the enrichment for genes associated with neuropsychiatric disorders, starting with 72 ASD-associated genes from large-scale exome analyses (false discovery rate (FDR) ≤ 0.001 equivalent to genome-wide significance, Fig. S7, S8, Table S1)[6]. Of note, considering a larger list of 255 ASD-associated genes at genome-wide significance (FDR ≤ 0.1) yielded similar patterns (Fig. S9).

Across the 29 clusters, corrected for neuronal fraction[8], all clusters showed some degree of enrichment (Table S2), reflecting the high expression levels of ASD-associated genes throughout the brain[39, 42]. Eight clusters were especially enriched for ASD genes, mapping to the thalamus, main olfactory bulb (MOB), and cortex (L2-3, L4-6-med) (Fig. 4). The highest GFES metric for ASD genes (0.89, 95% CI:0.88-0.90; Fig. 4a) was observed in the medial thalamus (TH-med) with the four other thalamic clusters (TH-endo, PF, TH-antdor, TH-mid) also in the top seven ASD enriched clusters by GFES. The neighboring, but functionally and anatomically distinct, epithalamus (EPI) showed relatively low enrichment (0.70, 95% CI:0.69-0.72; Fig. 4a). After thalamus, the main olfactory bulb was the most enriched, followed by two cortical clusters (L2-3, L4-6-med) with no significant difference between upper and lower cortical layers. Two other cortical layers (CTX-endo, L4-6-middle) showed relatively low enrichment.

**Figure 4:**
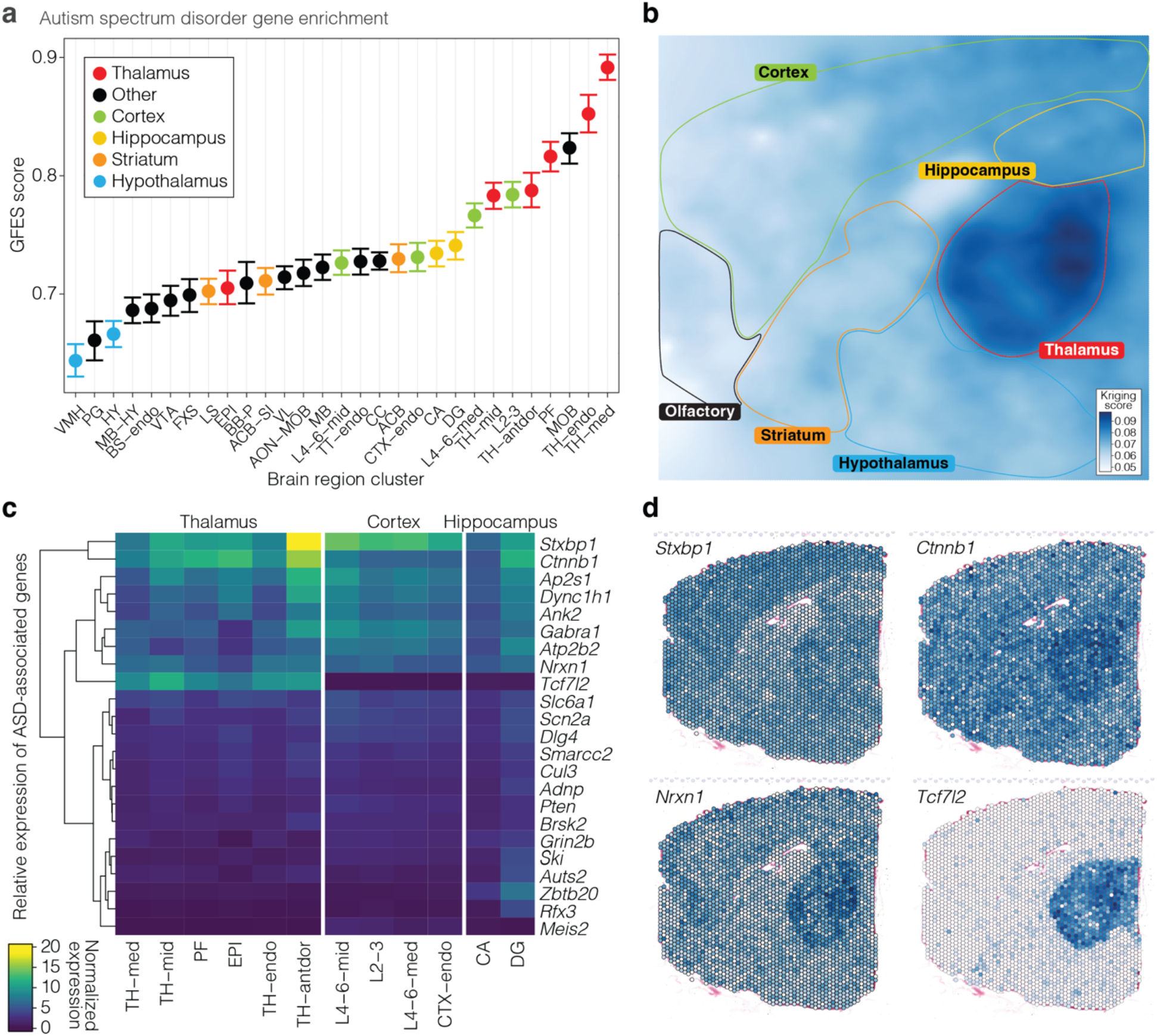
Enrichment of ASD-associated genes in the thalamus. **a)** GFES score based on 72 ASD-associated genes by regional cluster corrected for neuronal proportion. Error bars show 95% confidence intervals calculated by bootstrapping. b) Kriging plot showing spatial enrichment across all ASD genes assessed. **c)** Heatmap of gene expression across clusters for the most highly expressed genes. d) Spatial gene expression plots of specific ASD-associated genes.

The observed thalamic enrichment is especially marked for nine ASD-associated genes (Fig. 4c). The clearest thalamic pattern was for *Tcf7l2* (Fig. 4d), a transcription factor that is more highly expressed in neurons of the thalamus than the cortex in both mice and humans (Fig. S10[43]). Despite the strong *Tcf7l2* signal, thalamic enrichment of ASD-associated genes persists with its removal from the analysis (Fig. S9). Neuronal expression of *Nrxn1*, a pre-synaptic cell adhesion protein, and *Ctnnb1*, the Wnt signaling β-catenin protein show clear thalamic enrichment over other captured brain regions (Fig. 4d). Other ASD-associated genes are enriched in both the thalamus and cortex, such as *Stxbp1* which regulates the SNARE complex for neurotransmitter release.

### Schizophrenia genes are enriched in the olfactory bulb, cortex, hippocampus, and thalamus

Exome-based analyses identify ten genes associated with schizophrenia through rare protein-coding variation[9], but this is too few genes for reliable analysis of gene enrichment. We therefore used genes identified from common variation in a recent large-scale GWAS study, using the 435 reported high-confidence protein-coding genes in a “broader fine-mapped set” (Table S1)[10]. Like the ASD-associated genes, these are enriched for neuronal markers as shown previously[10] and replicated in our dataset (Fig. S7), so we applied the same correction for neuronal proportion as in ASD. Assessing spatial enrichment identified similar brain regions in schizophrenia and ASD (R^2^ = 0.37, p = 0.0005; see Fig. S11), with the main olfactory bulb (MOB), cortex (L2-3, L4-6-med), and thalamus (TH-med) being ranked highly by GFES in both disorders (Fig. 5a). Applying the Kriging geospatial model to both schizophrenia and ASD enrichment generated similar landscapes of gene enrichment (Fig. 5b, Fig. S11), although with a higher degree of enrichment in the dentate gyrus (DG) and Cornu Ammonis (CA) of the hippocampus in schizophrenia than ASD.

**Figure 5.**
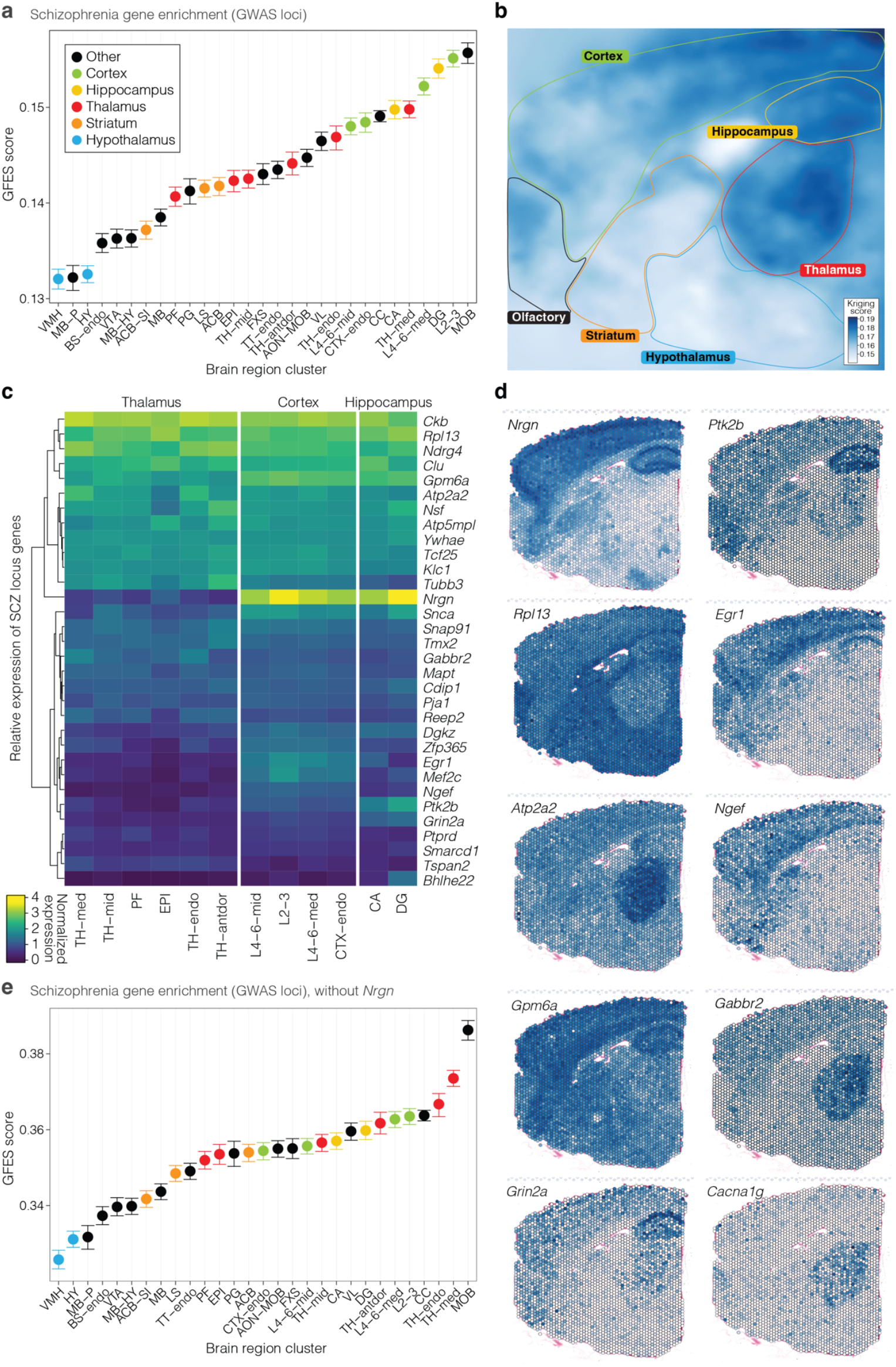
Enrichment of schizophrenia-associated genes in the olfactory bulb, cortex, thalamus, and hippocampus. **a)** GFES score based on 435 fine-mapped schizophrenia genes from GWAS by regional cluster corrected for neuronal proportion. Error bars show 95% confidence intervals calculated by bootstrapping. **b)** Kriging plot showing spatial enrichment across all schizophrenia genes assessed. **c)** Heatmap of gene expression across clusters for the most highly expressed genes. **d)** Spatial gene expression plots of key schizophrenia risk genes. e) Analysis in panel ‘a’ is repeated excluding the gene *Nrgn*.

As with ASD, a minority of the 435 genes contribute strongly to the regional enrichment patterns (Fig. 5c). Multiple schizophrenia fine mapped genes are enriched in the hippocampus and cortex, including neurogranin, *NRGN*/*Nrgn*, the tyrosine kinase *PTK2B*/*Ptk2b*, the ribosomal protein *RPL13/Rpl13*, and the transcription factor *EGR1/Egr1* (Fig. 5d).

However, within these four examples, the finemap posterior probability (FPP) values are low for *NRGN, PTK2B*, and *RPL13* (Table 1), raising the possibility that other genes might better reflect the causal genetic effects of the locus.[10] Of note, within the *NRGN* locus, another gene (*VSIG2*, a membrane bound immune regulator) has a higher FPP (0.38) than *NRGN* with the same index SNP[10], raising the possibility that *NRGN* does not contribute to schizophrenia liability. Removing *Nrgn*, the most prominent hippocampal gene, reorders the most enriched clusters, ranking the MOB and thalamus clusters at the top (Fig. 5e).

**Table 1.** Selected fine mapped GWAS loci for spatially dynamic genes from Trubetskoy *et al*.[10].

| Index SNP | Fine mapped SNP | Allele frequency | GWAS odds ratio | GWAS p-value | Finemap posterior probability | Human Gene | Human Ensembl ID | Mouse gene |
| --- | --- | --- | --- | --- | --- | --- | --- | --- |
| rs4766428 | rs4766428 | 0.46 | 0.93 | $2 \times 10^{-17}$ | 0.994 | ATP2A2 | ENSG00000174437 | Atp2a2 |
| rs778371 | rs778371 | 0.26 | 0.93 | $8 \times 10^{-17}$ | 0.546 | NGEF | ENSG00000066248 | Ngef |
| rs12498839 | rs12498839 | 0.07 | 0.89 | $1 \times 10^{-13}$ | 0.330 | GPM6A | ENSG00000150625 | Gpm6a |
| rs7915131 | rs7915131 | 0.43 | 1.04 | $4 \times 10^{-8}$ | 0.213 | ZNF365 | ENSG00000138311 | Zfp365 |
| rs1121296 | rs1121296 | 0.41 | 1.05 | $3 \times 10^{-9}$ | 0.202 | EGR1 | ENSG00000172260 | Egr1 |
| rs41533650 | rs41533650 | 0.20 | 0.93 | $8 \times 10^{-14}$ | 0.169 | GPM6A | ENSG00000150625 | Gpm6a |
| rs10985811 | rs10985811 | 0.21 | 0.95 | $3 \times 10^{-8}$ | 0.111 | GABBR2 | ENSG00000136928 | Gabbr2 |
| rs9926049 | rs9926049 | 0.27 | 0.95 | $3 \times 10^{-10}$ | 0.083 | GRIN2A | ENSG00000183454 | Grin2a |
| rs2022265 | rs2022265 | 0.47 | 1.05 | $3 \times 10^{-10}$ | 0.063 | SNAP91 | ENSG00000065609 | Snap91 |
| rs7938083 | rs7938083 | 0.29 | 0.95 | $2 \times 10^{-9}$ | 0.041 | TMX2 | ENSG00000213593 | Tmx2 |
| rs1940171 | rs55661361 | 0.32 | 1.06 | $1 \times 10^{-12}$ | 0.041 | NRGN | ENSG00000154146 | Nrgn |
| rs11076631 | rs12932628 | 0.41 | 0.96 | $4 \times 10^{-7}$ | 0.034 | RPL13 | ENSG00000167526 | Rpl13 |
| rs73229090 | rs2271920 | 0.37 | 0.96 | $1 \times 10^{-6}$ | 0.033 | PTK2B | ENSG00000120899 | Ptk2b |
| rs3752827 | rs3752827 | 0.33 | 1.05 | $5 \times 10^{-10}$ | 0.012 | YWHAE | ENSG00000108953 | Ywhae |

Focusing on the spatial dynamic genes with the higher fine-mapping estimates of the index SNP (Table 1) also supports the enrichment in the thalamus, cortex, and hippocampus. *ATP2A2*/*Atp2a2* is a calcium transporter associated with the endoplasmic reticulum and sarcoplasm and is enriched in the thalamus (TH-med). *NGEF/Ngef* is a neuronal guanine nucleotide exchange factor enriched in the cortex. *GPM6A/Gpm6a*, a neuronal membrane glycoprotein[44], enriched in both the dentate gyrus and thalamus and *GABBR2/Gabbr2* is a type B GABA receptor enriched in the thalamus (Fig. 5d).

Returning to the exome-derived list of ten genes, two have dynamic spatial profiles (Fig. 5d, Fig. S12). *GRIN2A*/*Grin2a* is enriched in the hippocampus, cortex, and thalamus and *CACNA1G*/*Cacna1g* is enriched in the thalamus (Fig. 5d). The gene *NRXN1*, enriched in the thalamus (Fig. 4d) is also robustly associated with both ASD and schizophrenia, mostly from copy number variants[45].

## Discussion

The contribution of specific brain regions to most human traits and disorders remains undefined, especially for neuropsychiatric disorders. Here, we investigate this question in novel spatial transcriptomic data from sagittal midsections of the adult mouse brain.

Site-specific transcriptomic data largely clustered by known anatomical regions (Fig. 2), such as the thalamus or cortex, with a few clusters defined by the intersection of anatomical regions and high proportions of endothelial marker genes (e.g., CTX-endo, TH-endo). To associate neuropsychiatric traits with these clusters, we developed GFES – the Gene Fraction Enrichment Score – to assess per spot gene enrichment controlling for covariates, such as cell type proportion. Because multiple prior analyses demonstrate that genes associated with neuropsychiatric traits are enriched in neurons[6, 8, 46], we used GFES to control for neuronal proportion focusing on the degree of gene enrichment by region rather than regional neuronal density.

Integrating hundreds of genes associated with ASD and schizophrenia with mouse spatial transcriptomic data using GFES, we assessed regional enrichment corrected for neuronal density. No single brain region emerged unequivocally in either disorder, instead multiple large brain regions were enriched. This fits with the observation that symptoms of ASD and schizophrenia are not a frequent consequence of individual brain lesions[47–49]. This may indicate that these disorders are not mediated by any individual brain region acting in isolation, but rather a circuit level phenomenon acting via neuronal impairment across multiple regions.

Nonetheless, some brain regions were substantially more enriched for ASD and/or schizophrenia genes than others. Regional enrichment was somewhat correlated between the two disorders (R^2^=0.37, Fig. S12) and similar regions ranked highly (thalamus, cortex, olfactory bulb) despite the ASD analysis using genes defined by exome sequencing and schizophrenia using GWAS and fine mapping. While clinically distinct, there is also epidemiological and genetic evidence of etiological overlap, including a five-fold increase schizophrenia relative risk in autistic individuals than in controls[50, 51] and shared genetic loci[6, 9].

The cortical enrichment in both conditions is consistent with the wealth of evidence linking this human-enlarged brain region to neuropsychiatric traits including transcriptomics, structural and functional neuroimaging, and the known role in cognition, behavior, and language[3, 15, 16, 52–56]. However, because the cortex is often analyzed in isolation, it is hard to contextualize this evidence. While our data support cortical involvement through the high enrichment values, in neither disorder is cortex the region most enriched for causal genetic factors.

The hippocampus has been assessed extensively as a model of synaptic plasticity[57], which, in turn, has been implicated in ASD and schizophrenia[58, 59]. However, there is less evidence directly linking the hippocampus to these traits[55, 56, 60, 61]. The hippocampus has a well-established role in memory[62, 63] and an emerging role in social behavior[64]. Our data support a role of the hippocampus in schizophrenia (Fig. 5), but this is less clear in ASD (Fig. 4).

Surprisingly, the strongest enrichment in ASD was observed in the medial thalamus (Fig. 4), a region also ranked fifth in schizophrenia (Fig. 5). Enrichment in the middle or anterior-dorsal thalamus was less pronounced; the limited availability of additional thalamic transcriptomic datasets prevents further resolution. Changes in the shape of the mediodorsal thalamus (included in the medial thalamus) and pulvinar thalamus (primate specific) are reported in ASD using structural MRI[65]. With functional MRI, overconnectivity between the cortex and thalamus have been described in both ASD and schizophrenia[3, 66–68] and a similar observation is observed in organoid models of the ASD- and schizophrenia-associated 22q11.2 microdeletion[69]. However, other analyses identify reduced thalamocortical connectivity in schizophrenia[70]. Our transcriptomic data are not able to define specific circuits; however, the enrichment in both the thalamus and cortex could be consistent with a role of thalamocortical circuitry. This thalamic enrichment for ASD-associated genes is consistent with analysis of human prenatal single cell data across development[71] and human prenatal spatial transcriptomic data[72].

The clearest thalamic-enriched ASD-associated gene was for *Tcf7l2*. In the human thalamus, *TCF7L2* is expressed in most cell types but especially excitatory neurons[73], whereas in the human cortex expression is primarily in progenitor cells and glia[74, 75] but not in mature neurons. *TCF7L2* is robustly associated with ASD (FDR = 0.0004[37]) and neurodevelopmental delay in exome-sequenced cohorts (FDR = 4 × 10^-11^)[37, 76] and characterized as clinical syndrome in a case series of 11 individuals[77]. *CTNNB1* follows a similar, but less dramatic expression pattern in human and mouse data[43, 73, 74] and similarly robustly associated with ASD (ASD FDR = 8 × 10^-5^, NDD FDR = 0). Of note, the proteins encoded by *TCF7L2* (TCF7L2, previously TCF-4) and *CTNNB1* (β-catenin) interact in the canonical Wnt signaling pathway[78]. β-catenin binds to TCF7L2 leading to transcriptional activation of Wnt target genes[79], so haploinsufficiency of either gene would be expected suppress Wnt target genes. To assess if the Wnt pathway is involved in ASD, we considered 254 genes in the Wnt signaling pathway, as defined by PANTHER Pathways (P00057) for gene ontology (GO:0016055). We observed a trend towards enrichment across the 72 most robust ASD-associated genes (OR: 3.31, 95% CI:0.88-Inf, p=0.07, Fisher’s exact test) that is significant with the more liberal 255 ASD-associated gene list (OR: 4.22, 95% CI:2.43-Inf, p=3.3×10^-5^, Fisher’s exact test). To date, a thalamic Wnt signaling pathway has not been defined.

We also observed substantial enrichment of the main olfactory bulb (MOB), which is the most enriched region in schizophrenia (Fig. 5) and third most enriched in ASD (Fig. 4). Olfaction is critical to social behavior and interaction in rodents, but it is hard to interpret this finding in humans where olfaction plays less of a role and the MOB is dramatically smaller[80]. Observed MOB enrichment may be indicative of the importance of sensory processing in neuropsychiatric traits and social behavior. Of note, assessment of postmortem gene expression in the cortex of individuals with ASD identified the most substantial dysregulation in sensory cortex (V1, visual cortex), rather than the prefrontal cortex[53].

We note several limitations to our analyses. First, assessments were based on the adult mouse, which may not capture regions that are transiently enriched in earlier developmental windows or human-specific expression patterns; notably analysis of human prenatal thalamus observes similar enrichment for ASD-genes[71, 72]. Future assessments would be enhanced by longitudinal data and integrated analysis with human data[46, 81, 82]. Second, while spatial transcriptomics enables systematic cross-regional analyses, the sagittal slices we selected exclude some major brain structures, such as the cerebellum, amygdala, and some hypothalamic nuclei. There is a need for more comprehensive multi-regional transcriptomic data. Third, we make the implicit assumption that high expression of genes associated with a trait indicates a causal role. This is logical for truly specific genes (e.g., *Tcf7l2* in the thalamus) but it is less evident for moderate quantitative differences; the sensitivity of GFES to highly specific genes is a strength in this context. Analyses including transcriptomic assessment of conditional heterozygous knockdown models of ASD/schizophrenia associated genes complemented by evidence of downstream disruption, e.g., protein levels, electrophysiology, and calcium imaging, will be necessary to validate these findings.

In conclusion, systematic analysis of multiregional spatial transcriptomic data supports the role of the thalamus and cortex in ASD and schizophrenia and also implicates the sensory structures and the hippocampus.

## Acknowledgements

We thank Mylinh Bernardi and Horng-Ru Lin of the Gladstone Genomics Core for their assistance with Visium spatial gene expression assay for fresh frozen tissue. Sequencing was performed at the UCSF CAT, supported by UCSF PBBR, RRP IMIA, and NIH 1S10OD028511-01 grants. This work was supported by a grant from the Simons Foundation Autism Research Initiative Sex Differences Collaboration (SFARI #736613 to S.J.S.), the National Institute of Mental Health (grant numbers R01MH125516 to S.J.S, R01MH129751 to S.J.S.), and HDRUK: Health Data Research UK QQ2 Molecules to Health Records Driver Programme to S.J.S.

## Materials and Methods

### Animals

The procedures of animal handling and tissue harvesting were approved by the Institutional Animal Care and Use Committee of University of California, San Francisco. All experiments were carried out on both male and female mice. The mouse strain (C57BL/6J) was obtained from The Jackson Laboratory and group housed in an UCSF barrier facility with a 12:12 hr light:dark cycle where food and water were available ad libitum.

### Sample preparation

We performed whole brain dissections on 8-week-old (P56) male and female mice and flash froze the samples in isopentane. The frozen samples were then embedded in O.C.T. (cryostat-embedding compound, Tissue-Tek, Torrance, CA) over dry ice to avoid cracking and stored at -80°C. We used a cryostat to collect 2 lateral and 2 medial sagittal sections of the brain of 14µm thickness, using anatomical markers to identify midsection slices just lateral to the midline. The lateral sections were approximately 100 - 150µm apart from the medial sections. We mounted 4 such sections from 1 brain on a Visium Spatial Gene Expression Slide (10x Genomics, 1000187) and stored at -80°C until further processing.

### Histology and sequencing

Spatial transcriptomics analysis was performed using the 10x Genomics Visium Spatial Gene Expression Assay. First, sections were placed on each of the four capture areas of the Visium Spatial Gene Expression slide. Each capture area contains a grid-like array of spots, where each spot contains primers to capture RNA lysed from permeabilized overlying tissue, including a unique spatial barcode of 12 bp to identify the spot’s location, a unique molecular identifier of 12 bp to distinguish each RNA transcript, and a poly(dT) sequence of 30 bp to capture mRNA for cDNA synthesis. After further cDNA amplification, the resulting libraries undergo Illumina sequencing, followed by alignment of transcript reads to the mouse genome using the 10x Spaceranger software package. Tissue fixation was performed in methanol at -20ºC for 30 minutes, then the slide was immediately stained with Hematoxylin and Eosin and imaged on an Automated Leica Aperio Versa 200 Slide Scanner at 20X magnification.

Next, permeabilization of tissue sections released the mRNA from cells, which were captured by primers on the Visium slide. Reverse transcription then added the spatial barcode, unique molecular identifier (UMI), and the partial Illumina sequencing Read 1 primer used for Visium. Following elution of samples from the Visium slide, one microliter of each sample was subjected to 25 cycles of qPCR. The optimal amplification cycles were determined using the Cq values at the exponential phase of the amplification plot, which is roughly 25% of the peak fluorescence. These values were used in the subsequent cDNA amplification reaction, followed by quality control on the Agilent Bioanalyzer. Ten microliters of cDNA were then used for fragmentation and library preparation, where dual indexes were added to the barcoded cDNA.

Quality control of the final libraries was completed on the Agilent 2100 Bioanalyzer using the High Sensitivity DNA Kit to determine the average library sizes, in addition to qPCR on the Applied Biosystems QuantStudio 5 Real-Time PCR System using the Roche KAPA Library Quantification Kit to determine the concentration of adapter-ligated libraries. Finally, libraries were pooled and sequenced on the NovaSeq 6000 S4 flow cell, paired-end 28×90 bp with 10 bp dual indexes.

### Spatial transcriptomics data processing

We processed the Visium sequencing data using the 10x Spaceranger software. To prepare the histology images, we opened whole-slide images in QuPath, rotated images by 180º, downsamples them to a longest edge size of 2000 pixels, and saved them as TIFF files. In a Docker container with Spaceranger installed, we converted FASTQ files to BAMs and raw matrices through the ‘spaceranger count’ pipeline, using default settings and referencing the ‘refdatagex-mm10-2020-A’ mouse transcriptome reference from 10x.

After generating raw matrices, we Seurat objects were created using default settings, which keeps only spots determined to be over tissue by Spaceranger’s image processing. For quality control, we plotted gene counts by UMI counts for each sample, log-scaling each axis. To identify objective cutoffs for each type of count, we clustered points using DBSCAN (Density-Based Spatial Clustering and Application with Noise) as implemented by the ‘dbscan’ package. To find the tuning parameter epsilon, the maximum distance between two data points to be considered neighbors, we located the “elbow” position at 0.1 in a K nearest neighbor distance plot. For the minimum points parameter, we used a standard value of 3 based on 2 dimensions + 1. Clustering separated each sample into two dominant groups, with one large, dense cluster and a smaller, looser cluster. Clustering all spots from all samples together replicated this pattern and identified a threshold of about 100 UMIs and 100 genes between the clusters, which we took as the UMI and gene lower cutoffs along with a 30% mitochondrial upper cutoff for low-quality spots.

Once we filtered spots, we normalized batch effects by using the Seurat SCTransform data integration pipeline. After separately normalizing each sample batch (slides 1-2 and 3-4) by SCTransform, we performed anchor-based integration across batches using default settings except increasing the number of features to 3,000 for greater resolution.

### Clustering and marker identification

We clustered spots using the Louvain-based shared nearest neighbor modularity optimization method implemented in Seurat, with default settings except increasing the number of dimensions to 30 when finding neighbors and reducing the resolution to 0.5. To visualized clusters, we plotted them after Uniform Manifold Approximation and Projection (UMAP) dimensional reduction, also generated with dimensions increased to 30.

To identify tissue specific gene markers, we first found differential markers using Seurat, comparing each cluster against all other clusters using default settings except increasing the minimum percentage threshold to 25% to find more broadly expressed genes. To refine this list, we took the markers with the greatest difference in percent expression within versus outside of the cluster and further filtered by the top log of the ratio of these percentages. Next, we applied the tau metric[32] as implemented in Kryuchko-Mostacci & Robinson-Rechavi[31] to each gene as a measure of tissue specificity to find the genes with the most focal, cluster-specific expression[83].

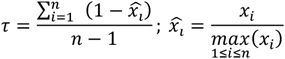

### Atlas alignment

To identify clusters anatomically, we mapped our samples into a common coordinate reference space using the Allen CCFv3 3D atlas to perform segmentation by registration. First, we manually identified the closest sagittal atlas slice corresponding to each of our sections. Next, we performed image registration to move each atlas section into alignment with the corresponding section. We used the Elastix image registration toolkit as implemented in our MagellanMapper software application[25]. After rescaling, rotating, and flipping the atlas section to approximate the sample orientation to aid registration initialization, we performed rigid registration using 1024 iterations of translation and 0-4096 iterations of affine transformations depending on the necessary cycles to align images visually. After rigid registration, we added deformable registration using 0-256 iterations of b-spline transformation with a voxel size of 150.

While these standard transformations typically aligned external borders well, internal structures often remained slightly misaligned, likely because of the multimodal registration[84]. To guide internal structure alignment in a semi-automated fashion, we identified clusters that clearly corresponded to large atlas regions, such as thalamic and hippocampal clusters. Using the corresponding-point b-spline transformation with 256 iterations and a voxel size of 30-50, we aligned the centers of each specified cluster-region pair. This atlas alignment allowed us to segment the samples using the atlas region annotations, where we assigned each spot to the corresponding atlas label at the spot’s center pixel. To visualize the regions that mapped best to each cluster, we measured the proportion of spots in each cluster assigned to each atlas region and plotted these proportions as a heat map.

### Cell type and region deconvolution

Each spot represents an amalgam of neighboring cells, potentially from diverse cell types. To estimate the proportion of cell types in each spot, we performed deconvolution using Robust Cell Type Decomposition (RCTD) from the spacexr package[85]. We ran each deconvolution in “full” doublet mode for no restrictions on cell type number and normalized the cell type proportions to a total of one per spot. To visualize these proportions, we plotted each cell type weighting on each sample to assess for expected locations. We also aggregated the mean weightings for each cell type within each cluster as a heat map. For region-specific datasets, such as the HypoMap hypothalamic atlas, we first subset our dataset to the corresponding region based on the registered anatomical delineations before performing the deconvolution.

### Gene enrichment

We computed the gene enrichment fraction at each spot using the equation:

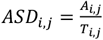

where *A*_*ij*,(_ = ASD gene UMIs at spot *i* in section *j, T*_*ij*,(_ = total gene UMIs at the same spot, making *ASD*_*ij*,(_ is the gene enrichment fraction.

To measure and compare the gene enrichment by cluster, we next performed an ANCOVA test as implemented in the R ‘aov’, where gene enrichment is the dependent variable, and cell type proportion is the categorical independent variable. Plots of the enrichment fraction versus neuronal proportion for each spot revealed heteroscedasticity — increased variance as neuronal fraction increased — which stabilized after applying a square root transformation of the enrichment values. We also observed a slight quadratic effect, which we modeled with a polynomial, resulting in the ANCOVA formula: 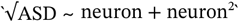 .We took the enrichment of each cluster to be the estimates as output from ‘summary.lm’.

To identify statistical differences among regions, we could not use the estimates’ p-values because the spots are spatially correlated with one another. Instead, we applied a bootstrapping resampling technique to measure 95% confidence intervals. For each cluster, we resampled its spots with replacement and reran the ACOVA across all clusters, generating a new set of estimates. We repeated this process 1000 times, resulting in an estimate of the sampling distribution of ANCOVA estimates and their 95% confidence intervals.

To further validate the gene enrichment metric, we tested its ability to identify enrichment in simulations of gene expression. We simulated an RNA-seq dataset with 20,000 cells and 3,000 genes, where the number of UMIs for each gene in each cell was randomly taken from a uniform distribution. To simulate a baseline number of “expressed” genes, we randomly sampled 10% of all cell-gene pairs to have three times their original UMIs. To test whether the metric could detect differences in enriched clusters, we further expressed 10% of genes in two clusters, which demonstrated clear enrichment compared to the other clusters using the metric (Fig. S6a).

We also tested subtler enrichment by linearly increasing the number of expressed genes in the cells of three clusters. As expected, the metric showed a linear increase in enrichment across these clusters (Fig. S6b). To test non-linear enrichment, we increased the number of expressed genes exponentially across the whole simulated dataset, leading to an exponential enrichment curve using the metric, as anticipated (Fig. S6c).

In some cases, a single strongly expressed gene dominated the final enrichment results. To identify such genes, we inspected heatmaps of gene expression by cluster and compared results with and without these genes. We also compared enrichment on raw UMI counts versus normalized counts.

### Geospatial interpolation through Kriging

To model gene enrichment spatially, we applying the Kriging geospatial interpolation technique[86] implemented in the ‘gstat’ package to predict values at higher resolution while accounting for covariates. Kriging models spatial autocorrelation, where points that are closer together are expected to be more similar. To model this autocorrelation, we first generated a semivariogram, which plots the variance of pairs of points by their distance, binned into similar distances. Next, we fit the variogram by empirically determining the sill (variance when the model starts to flatten), range (distance at this point), nugget (variance when distance is 0), and the mathematical model. With our fit variogram model, we used Ordinary Kriging to map it to a dense grid of points over each sample to predict the gene enrichment at each point using a weighted average of nearest neighboring points. To incorporate trends from the neuronal proportions, we applied Universal Kriging with neurons as the covariate, using the same formula as for the ANCOVA.

## Software versions

We used the following software versions: Space Ranger 2.0, QuPath 0.3.2, Seurat 4.1, dbscan 1.1, Elastix 5.0, MagellanMapper 1.6, spacexr 2.0, gstat 2.1, R 4.2, Python 3.8, Java 17.

## Data availability

The data will be released upon publication. The raw, unprocessed spatial transcriptomics data will be available on GEOXXX as FASTQ files. The processed data will be available as an RDS Seurat object file, also on GEOXXX.

## Code availability

The source code including the enrichment pipeline will be released upon publication on GitHub at XXX.

