## Supplemental Figures for "Spatial transcriptomics implicates the thalamus and cortex in autism and schizophrenia"

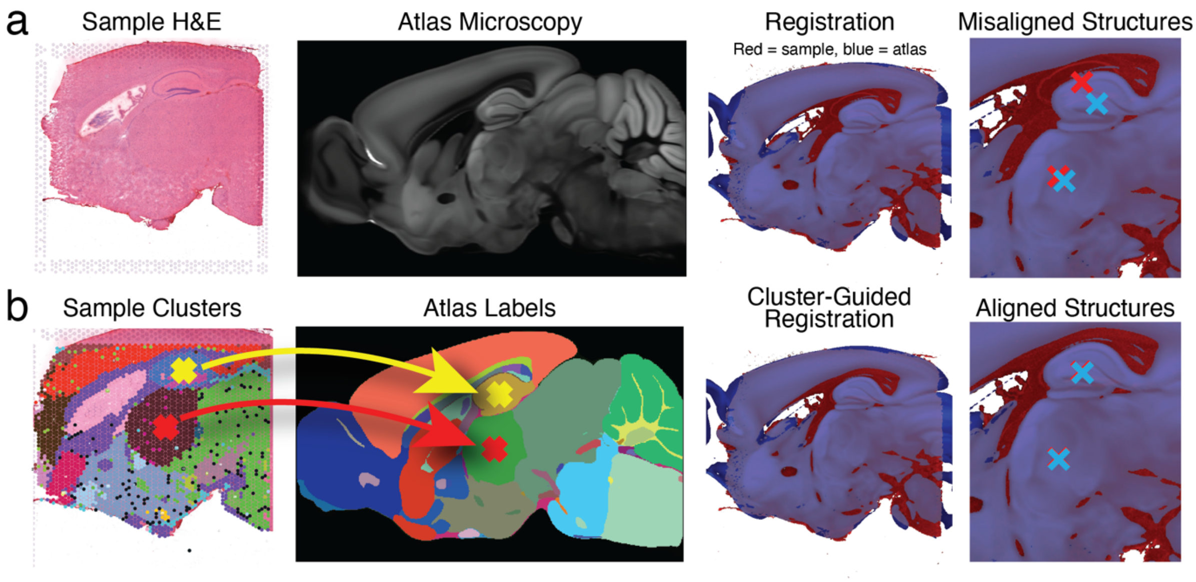


**Figure S1.** Cluster-guided atlas image registration. (a) Atlas image alignment. To segment the sample H&E image by region (far left), the CCFv3 atlas intensity image (middle left) is moved into alignment with the sample, as shown in the overlaid registration (middle right). Although the outer borders align reasonably well, closer inspection of the internal structures such as the hippocampus (far right, top x's) and thalamus (bottom x's). (b) Atlas alignment refinement. Clearly identified transcriptomic clusters (far left) corresponding to atlas labels (middle left) are used to guide the final step of the transformation (middle right), tightening up alignment in internal structures while preserving outer border alignment.

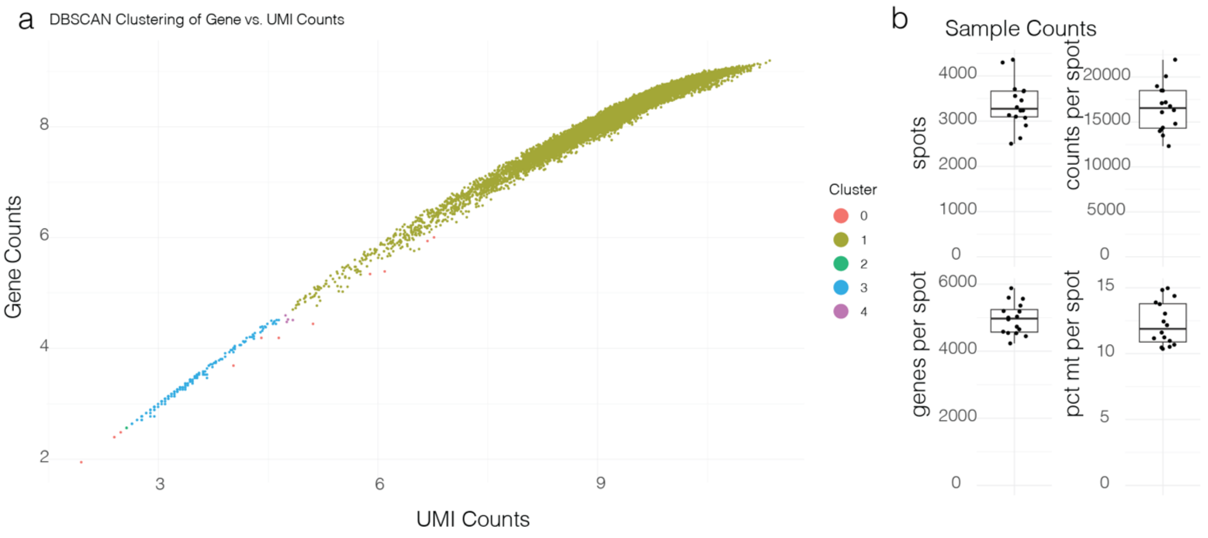


**Figure S2.** Quality control metrics. (a) Clustering by DBSCAN of all spots' gene number by UMI counts. Spots clustered into two dominant groups (blue and gold), demarcating cutoff thresholds for filtering lower-quality spots. (b) Box plots of total spots (upper left), UMI counts per spot (upper right), number of unique genes per spot (bottom left), and percent of mitochondria transcripts per spot (bottom right).

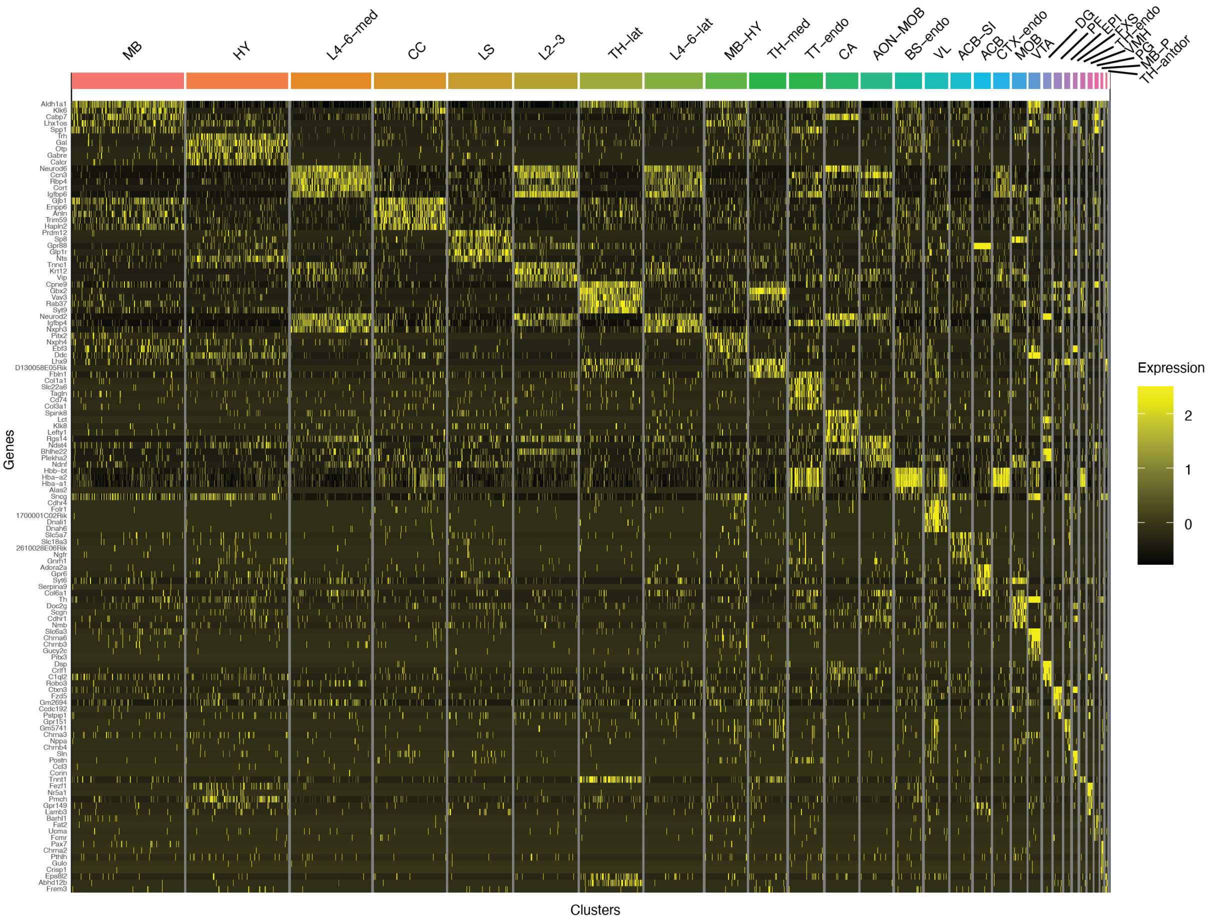


**Figure S3.** Differentially expressed markers filtered by tissue specificity. Gene expression of the top five differentially expressed markers for each cluster are shown across all clusters.

**a** Gene Enrichment Fractions By Neuronal Proportion

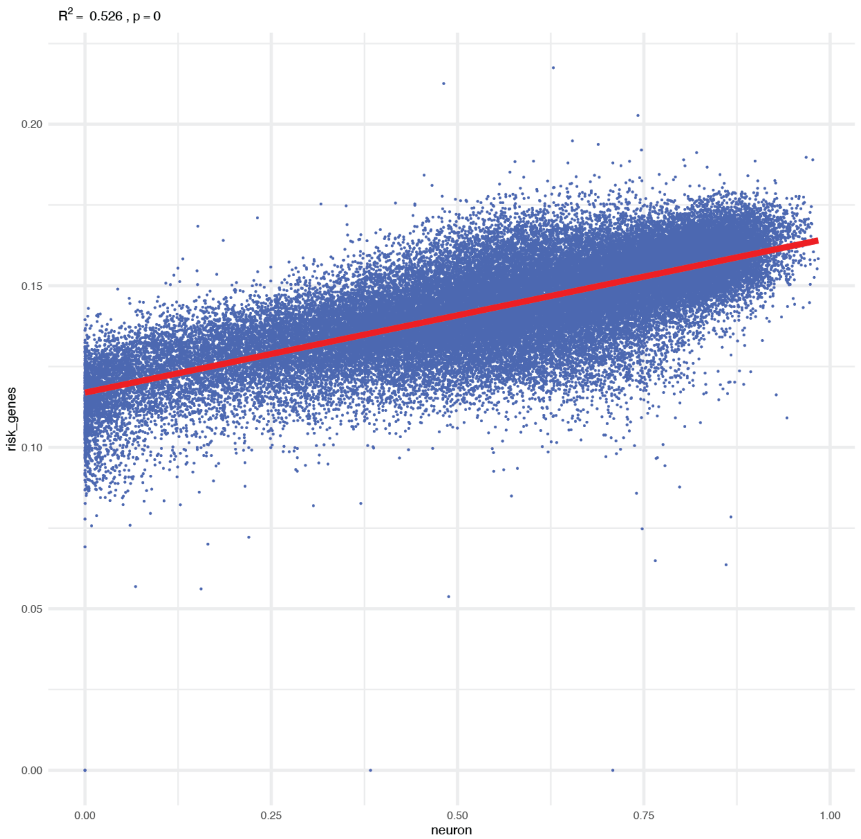


**b** Enrichment By Neurons For Cell Types

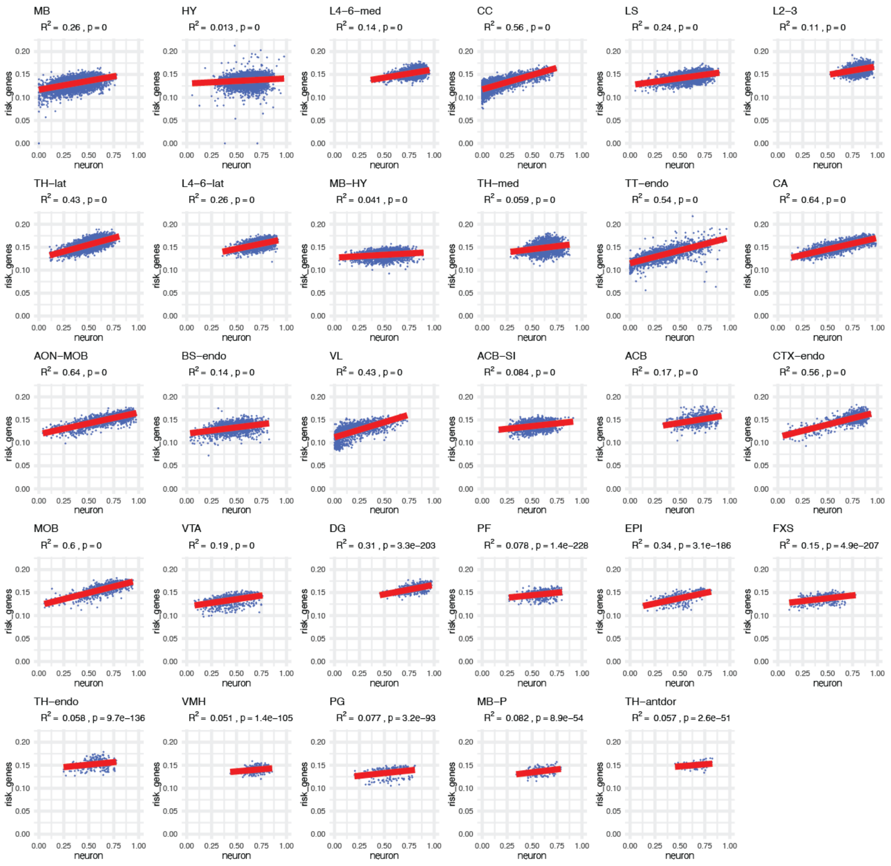

**Figure S4.** Correlation of gene enrichment with neuronal proportion. (a) Correlation of ASD risk gene enrichment fractions for each spot by predicted neuronal proportion within each spot across the full dataset (left) and split by cluster (right). Gene enrichment fractions were transformed by a square root to account for heterskedasticity, while neuronal proportions were modeled by a polynomial to account for a small quadratic effect. (b) Correlation of height and (c) schizophrenia gene enrichment fractions by neurons across all spots.

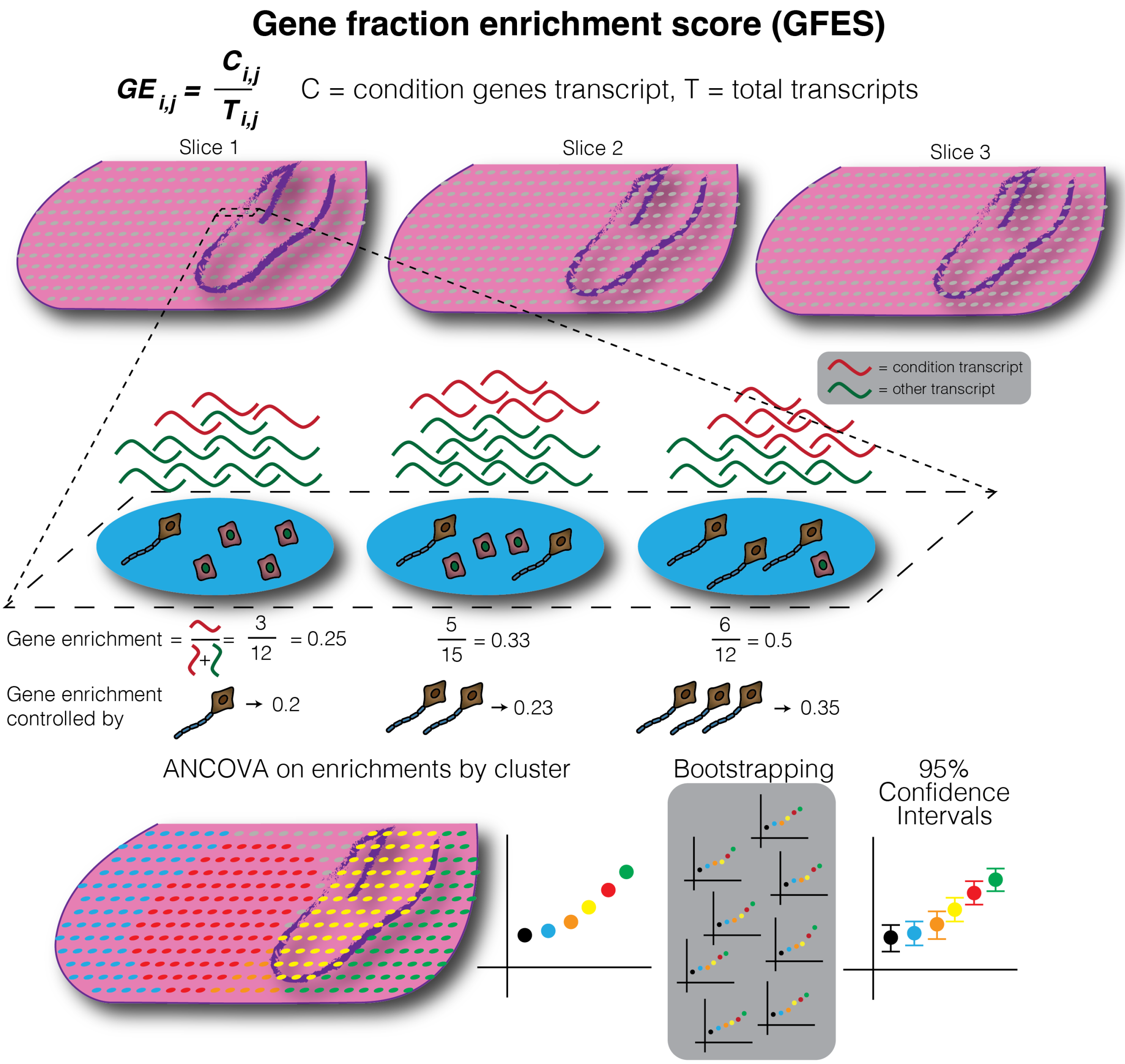


**Figure S5.** Schematic of the gene enrichment fraction score (GEFS) metric. Gene enrichment (GE) by spot or cell is taken as a fraction of the condition genes' UMI transcripts (red) over the total transcripts (red plus green). GE's are grouped by cluster, and each cluster's total enrichment is compared using ANCOVA. A bootstrapping resampling technique provides 95% confidence intervals to show statistically significant differences among clusters. To account for differences in spot characteristics such as cell type composition, GE's can be controlled for cell type such as neuronal proportion by incorporating it as a covariate in the ANCOVA statistic.

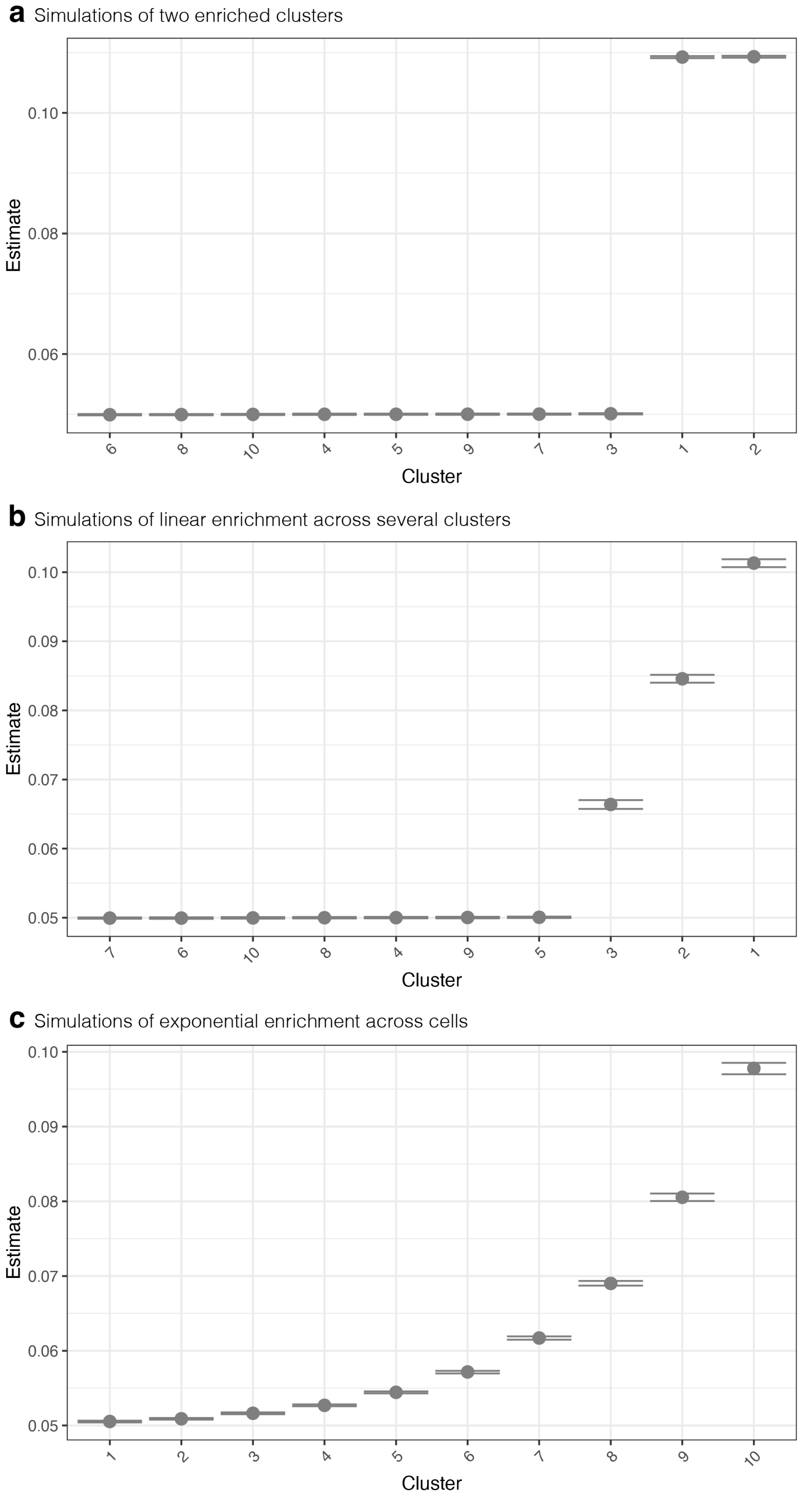


**Figure S6.** Gene enrichment analysis of simulated gene expression. (a) Simulated elevation in the gene expression of 10% of the genes in clusters 1-2 yielded the expected selective enrichment in these clusters as measured by GFES. (b) A linear increase in the number of genes with elevated expression across clusters 1-3 were detected as a corresponding linear increase in enrichment. (c) Exponential increase in number of genes with elevated expression across all cluster resulted in an exponential enrichment pattern.

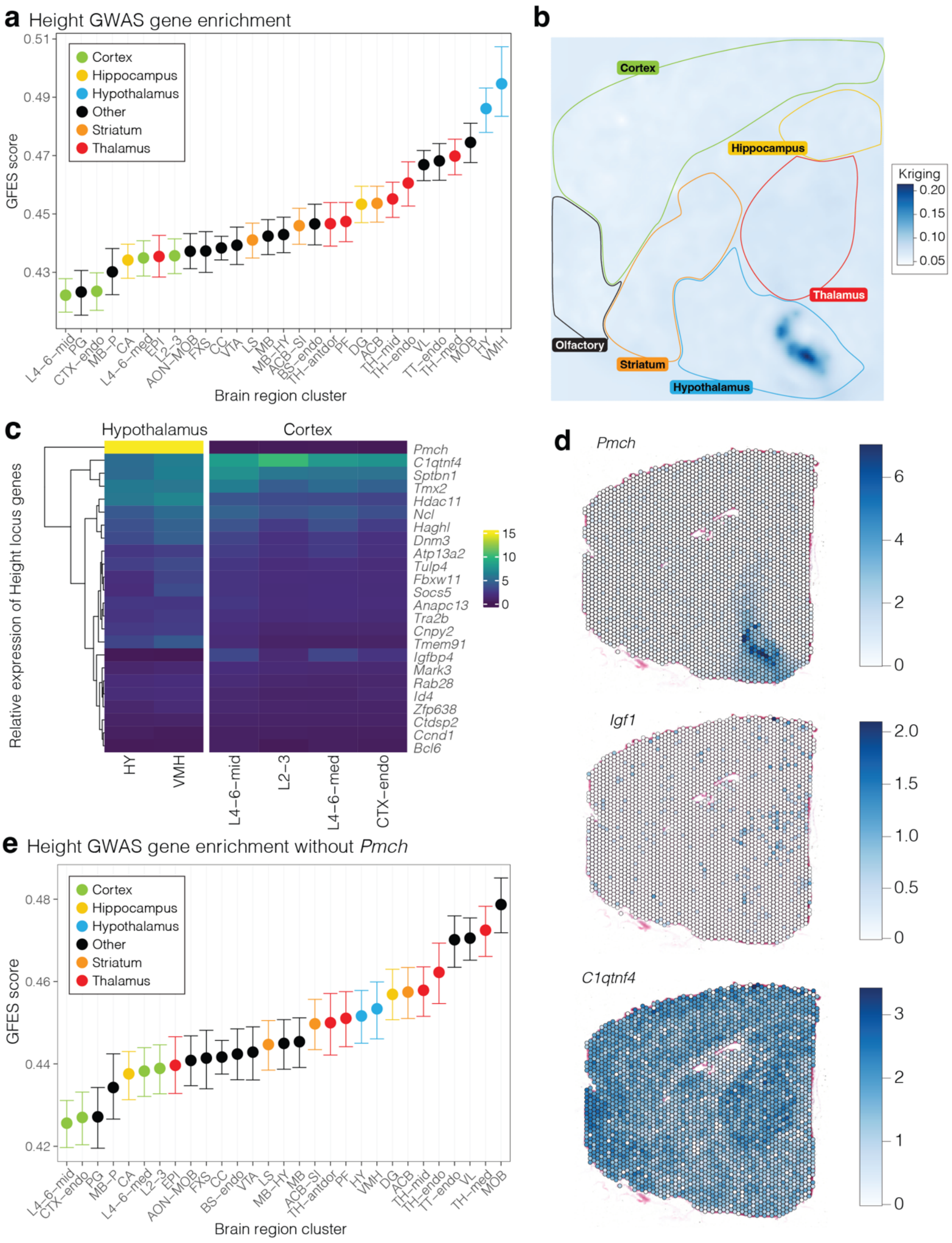


**Figure S7. Height enrichment corrected by neuronal proportion. a)** GFES score based on 188 genes associated with height from GWAS by regional cluster corrected for neuronal proportion. Error bars show 95% confidence intervals calculated by bootstrapping. **b)** Kriging plot showing spatial enrichment across all height genes assessed. **c)** Heatmap of gene expression across clusters for the most highly expressed genes. **d)** Spatial gene expression plots of key height genes. **e)** Analysis in panel 'a' is repeated excluding the gene *Pmch*.

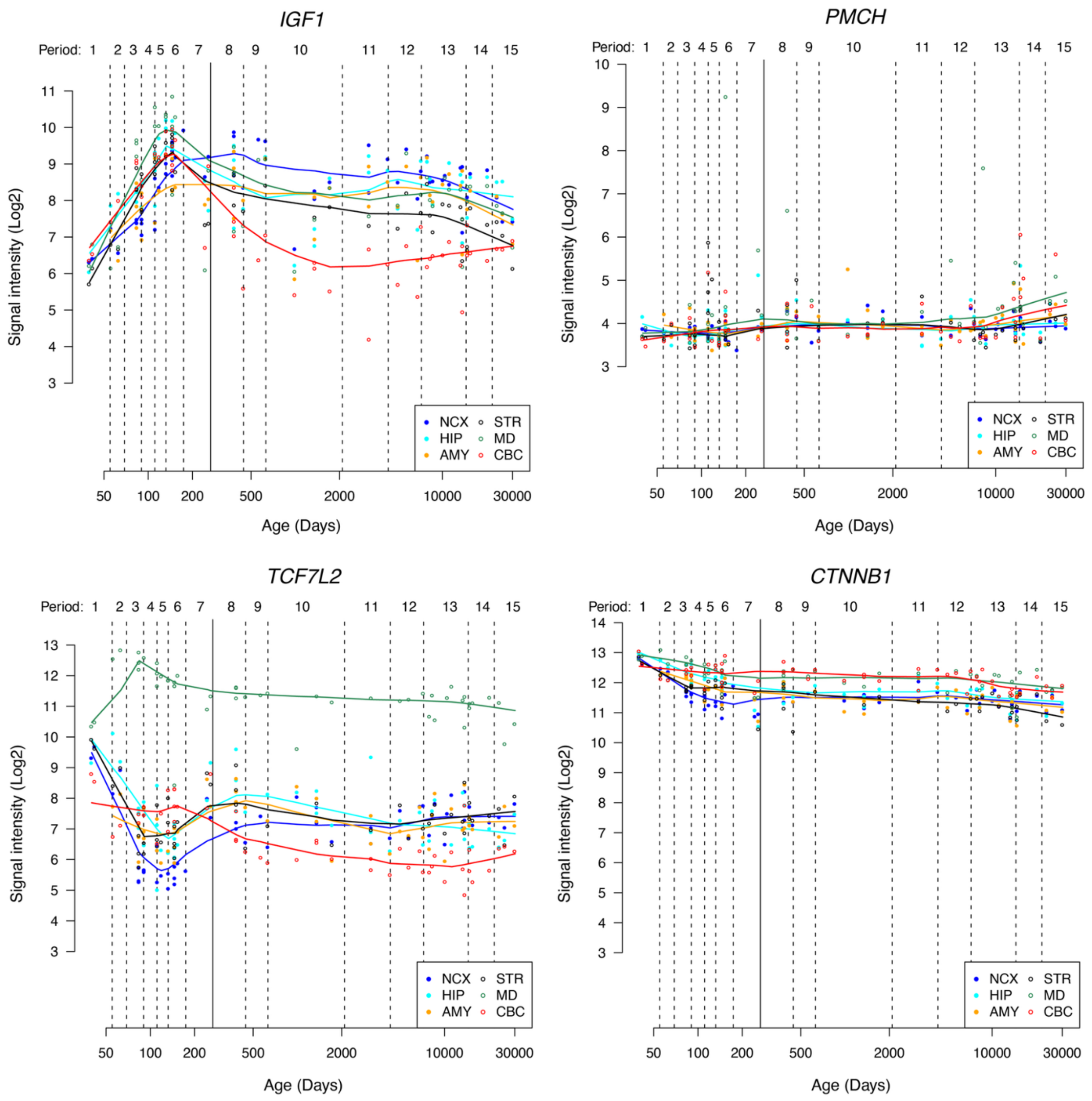


**Figure S8. Expression of genes in the developing human brain.** Gene expression is shown for four genes (*IGF1, PMCH, TCF7L2, CTNNB1*) in six brain regions (neocortex [NCX], hippocampus [HIP], amygdala [AMY], striatum [STR], mediodorsal thalamus [MD], and cerebellum [CBC]) across development in the human brain. Each point represents on same. The x-axis shows log-scaled postconceptual days with developmental periods bound by dashed lines. Data and period definitions are from Kang et al. 2011 (PMID: 22031440).

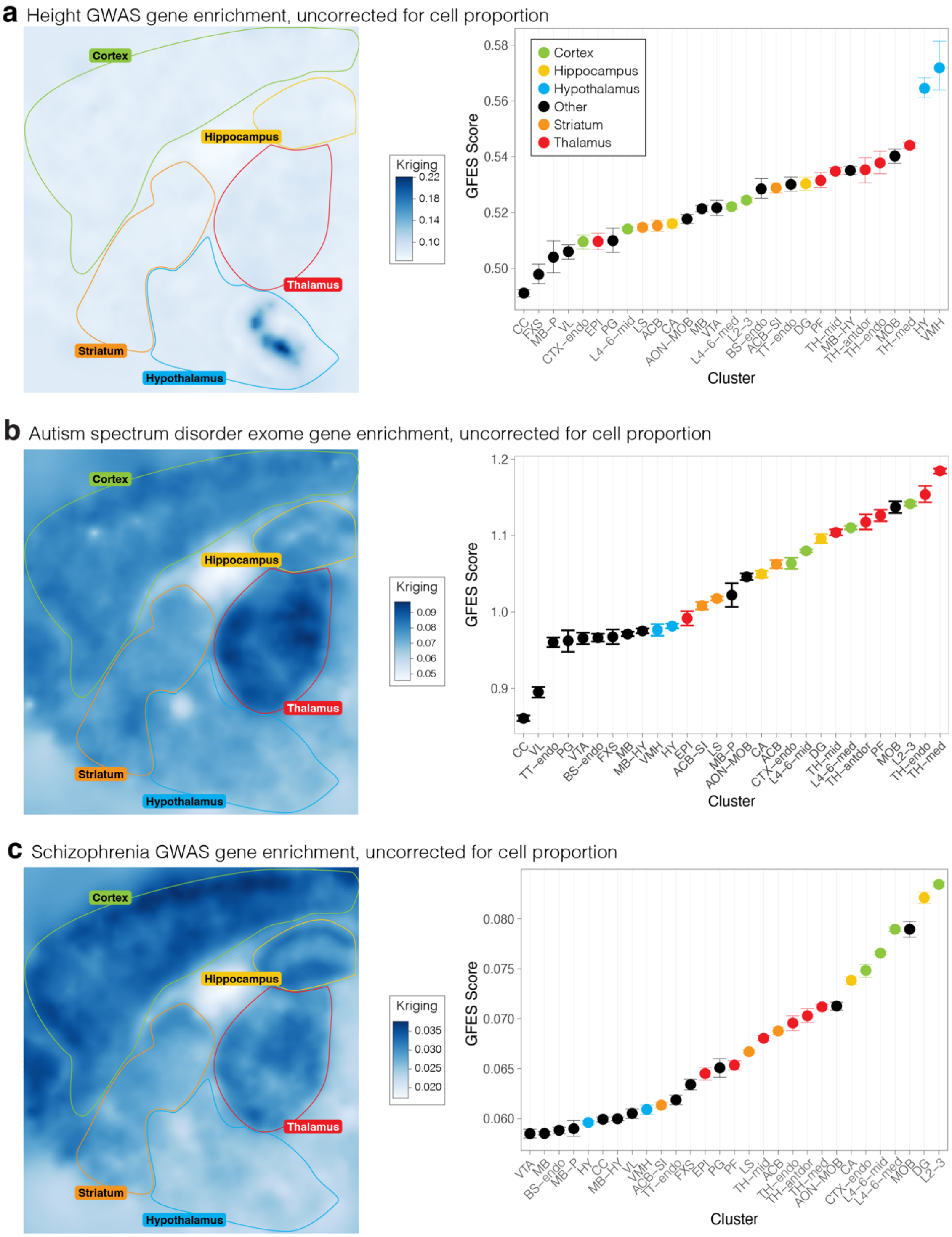


**Figure S9. Gene enrichment uncorrected for neuronal proportion. a)** Kriging plot showing spatial enrichment across 188 genes associated with height from GWAS without including neuronal proportion as a co-variate (left). GFES scores are shown for the same data (right); error bars show 95% confidence intervals calculated by bootstrapping. **b)** Equivalent data for 72 ASD-associated genes. **c)** Equivalent data for 435 schizophrenia-associated genes.

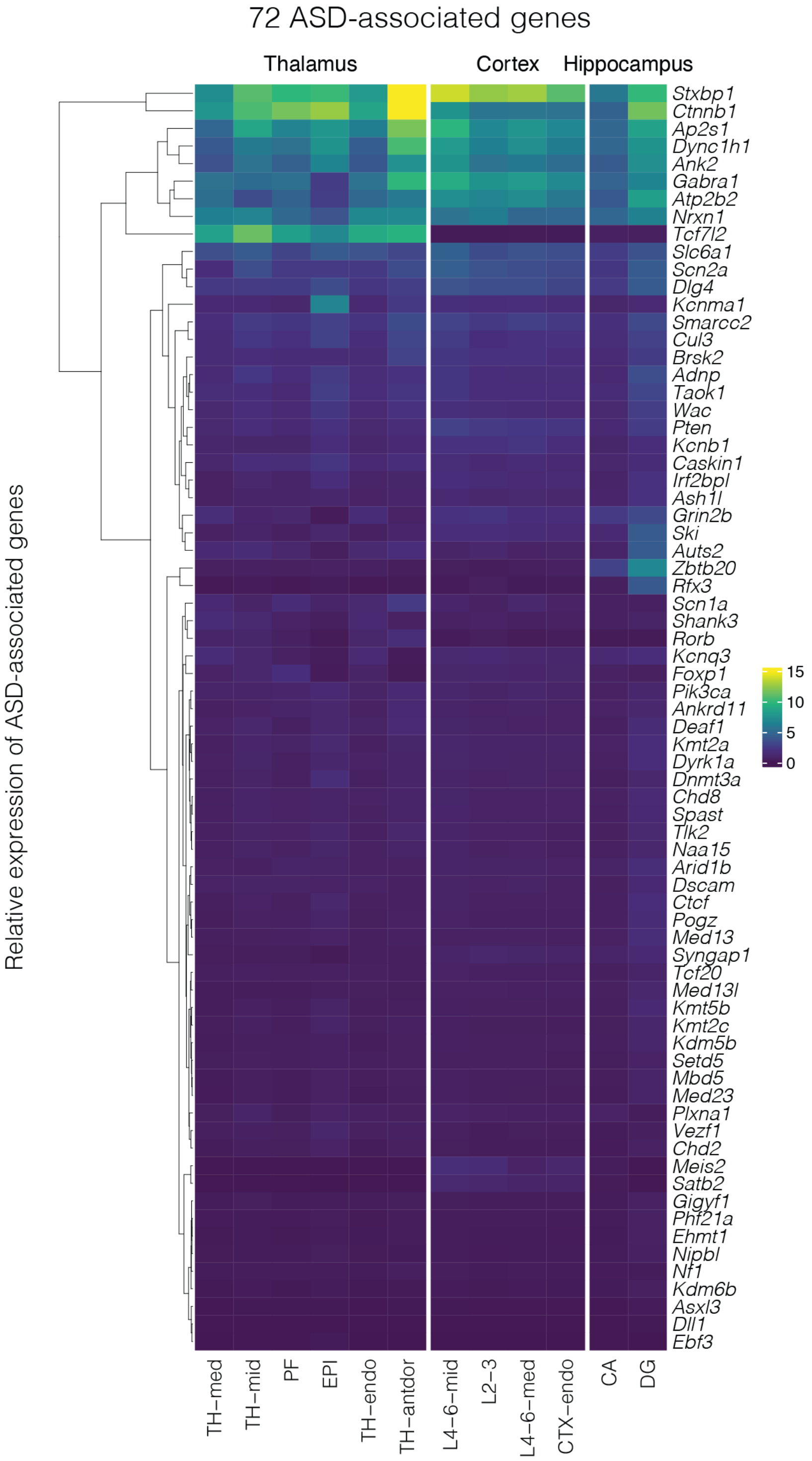


**Figure S10.** Relative expression across ASD-associated genes. Heatmap of gene expression by clusters within the thalamus, cortex, and hippocampus for the most strongly ASD-associated genes.

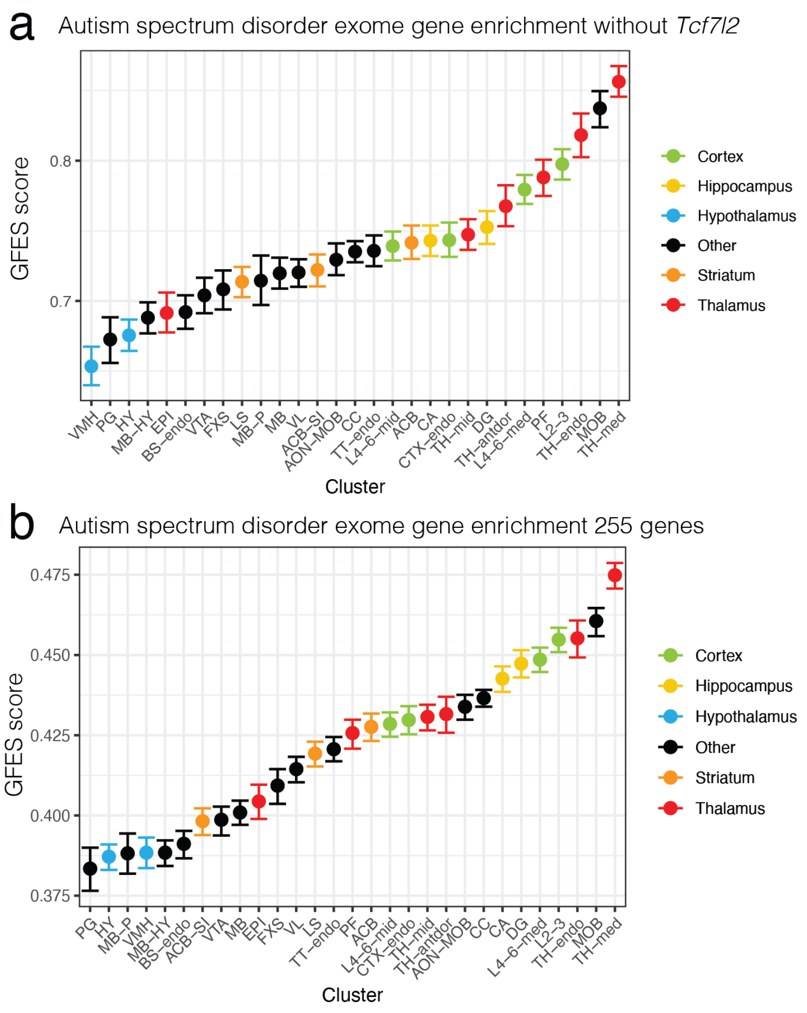


**Figure S11. Relative expression across ASD-associated genes. a)** GFES scores are shown for 71 ASD-associated genes, having excluded *TCF7L2* (which showed the greatest degree of thalamic enrichment); error bars show 95% confidence intervals calculated by bootstrapping. **b)** Equivalent plot for 255 ASD-associated genes based on a more permissive FDR threshold (0.1).

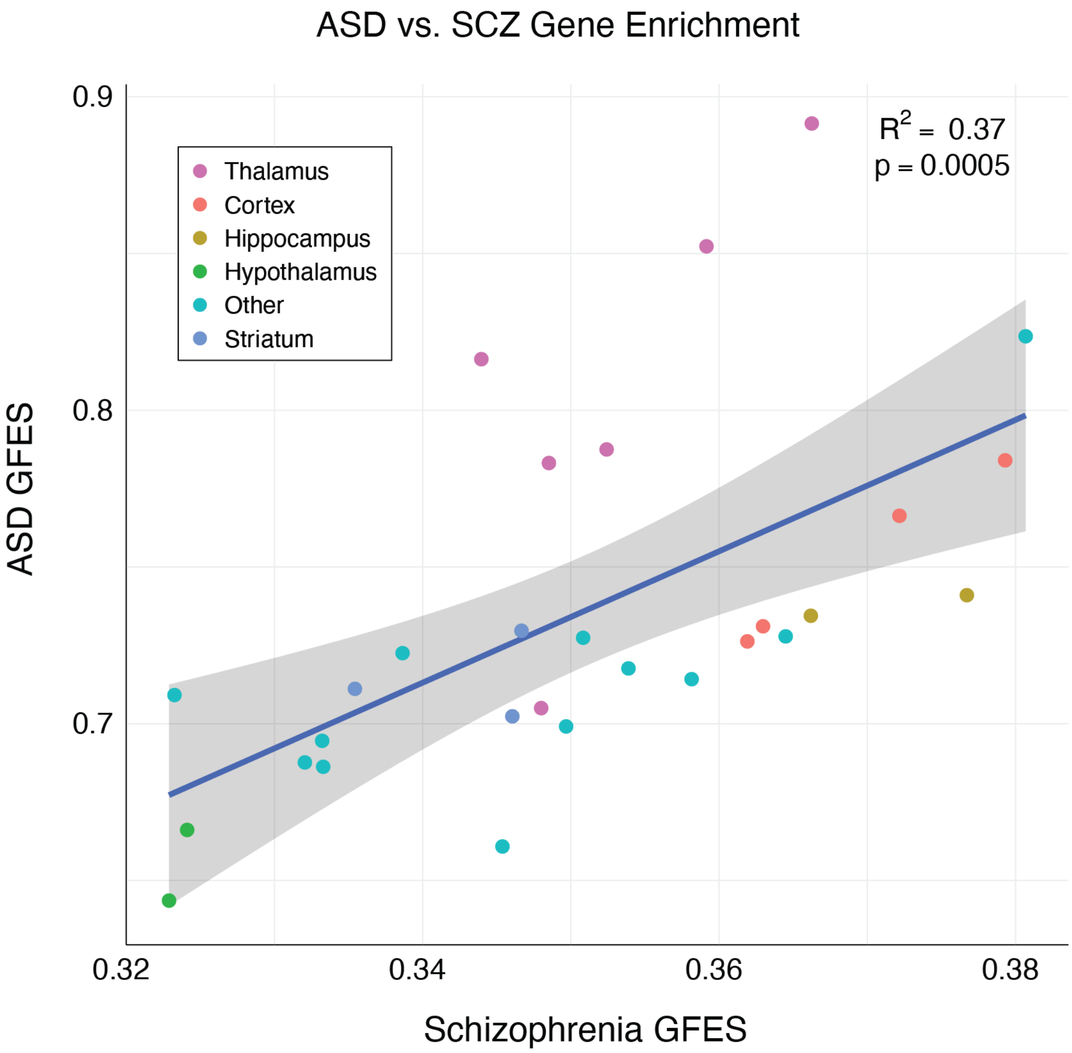


**Figure S12. Enrichment by cluster between ASD and Schizophrenia.** GFES scores are shown for each cluster from 72 ASD-associated genes and 435 schizophrenia-associated genes. Color indicates cluster super-regions.

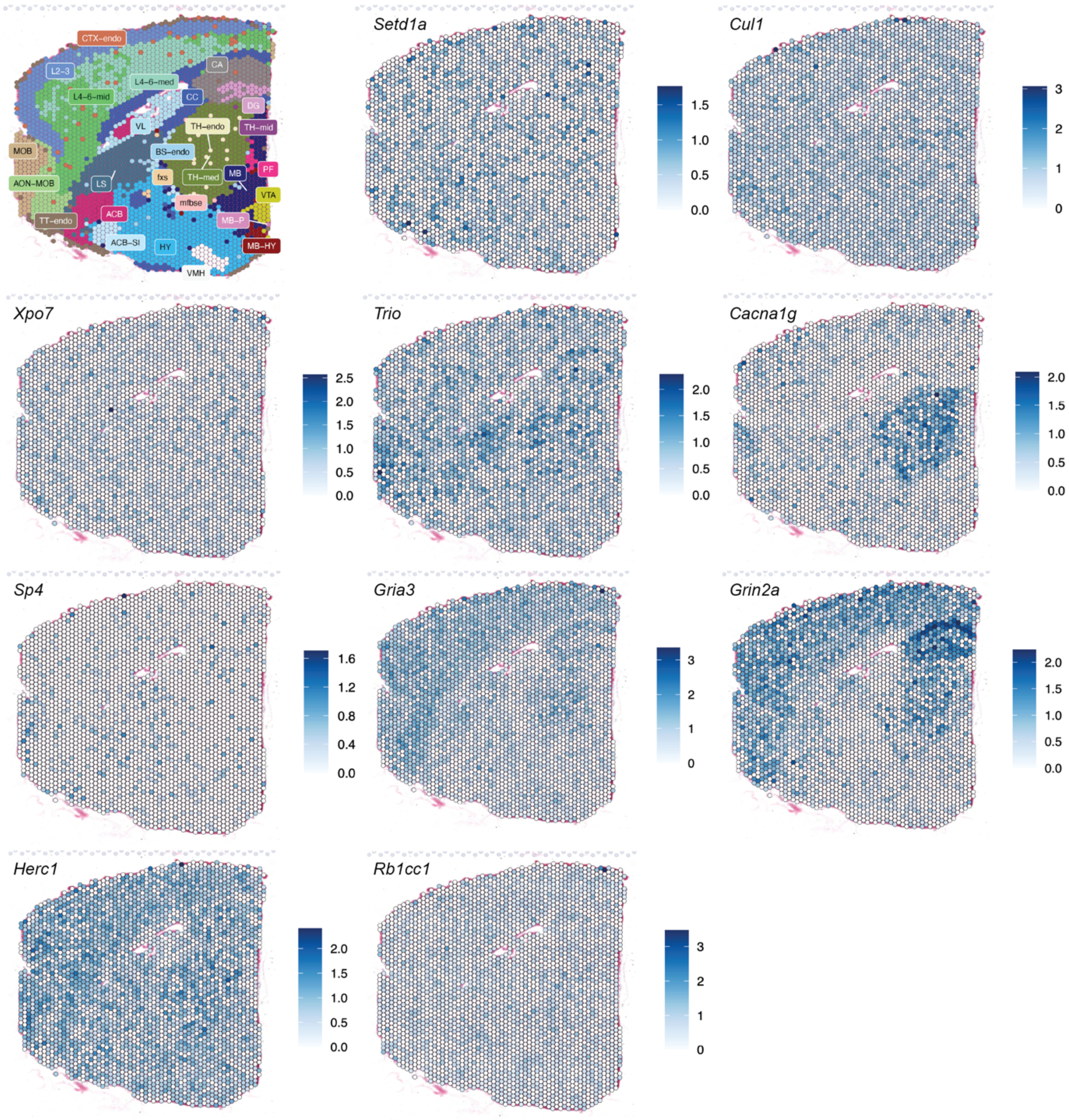


**Figure S13. Expression patterns of schizophrenia-associated genes from exome sequencing.** Spatial gene expression plots of ten schizophrenia genes identified by exome sequencing (PMID: 35396579).
